# Proliferative Capacity and Neural Lineage Commitment of Müller Glia in the Adult Human Retina

**DOI:** 10.64898/2026.04.27.717633

**Authors:** Dániel Péter. Magda, Teadora Tyler, Lili Gerendás, Ferenc Kilin, Anita Csorba, Bence György, Simone Picelli, Zoltán Zsolt Nagy, Arnold Szabó

## Abstract

**Background:** The mammalian retina lacks meaningful regenerative capacity, and degeneration usually leads to irreversible vision loss. Although lower vertebrates regenerate retinal neurons through Müller glia, this capacity has generally been considered absent in humans. This study investigated whether defined humoral cues alone are sufficient to unlock a latent neurogenic program in human Müller cells.

**Methods:** Long-term organotypic retinal cultures were established from 39 adult donors. Cultures were treated with FGF-2 and GSK-3 inhibition to assess proliferation across the peripheral and central retina. Cellular responses were evaluated using single-cell transcriptomics and immunohistochemistry.

**Results:** FGF-2 treatment and GSK-3 inhibition induced robust proliferation across both peripheral and central retina, with 79.09 ± 6.32% of dividing cells identified as Müller glia, some completing multiple cell cycles. Single-cell transcriptomics revealed activation of progenitor-like and neuronal differentiation pathways, whereas immunohistochemistry demonstrated expression of early and late neuronal markers spanning all major retinal lineages. Newly generated cells expressed markers of cone, rod, bipolar, horizontal, amacrine, and ganglion cell identities, together with evidence of early synaptogenesis.

**Conclusions:** These findings reveal an intrinsic regenerative potential in adult human Müller glia and show that defined humoral cues can activate a latent neurogenic program in these cells. This may have implications for future vision-restoration strategies in degenerative retinal disease.

## Background

The mammalian retina is a highly specialized neural tissue responsible for detecting light and initiating visual processing. Vision loss resulting from the degeneration of retinal neurons—such as photoreceptors and retinal ganglion cells—in diseases including age-related macular degeneration, glaucoma, diabetic retinopathy, and retinitis pigmentosa represents a major global public health challenge(1, 2). These conditions are characterized by the progressive and often irreversible loss of neurons and their intricate synaptic circuitry(3, 4). Current therapeutic strategies offer no definitive cure, as none can fully restore the native architecture or function of a healthy retina. Consequently, regenerative medicine has emerged as a promising frontier, aiming to replenish lost neuronal populations and reconstruct visual circuitry by leveraging either transplanted cells or endogenous repair mechanisms(5, 6). Of particular interest is the activation of endogenous repair pathways, particularly those mediated by Müller glia—the principal glial cells of the retina—which represent a promising therapeutic avenue(7, 8).

Unlike mammals, certain non-mammalian vertebrates, such as zebrafish, possess a remarkable capacity for retinal regeneration(9). In these species, Müller glia function as intrinsic stem-like cells that, upon injury, de-differentiate, proliferate, and give rise to progenitor cells capable of regenerating all major retinal neuron types(10). This regenerative response is orchestrated by a tightly regulated cascade of developmental signals, including the induction of transcription factors such as Achaete-scute homolog 1 (ASCL1) and the activation of key signaling pathways such as Notch, Wnt/β-catenin, and Jak/Stat(11, 12).

In contrast, mammalian Müller cells respond to retinal injury predominantly through reactive gliosis, characterized by inflammation and scar formation, rather than neurogenesis(13). The absence of a regenerative response is associated with a failure to activate key proneural genes, such as ASCL1, which remains silent in the adult mammalian retina(14). However, studies in rodents have demonstrated that, under defined experimental conditions, Müller cells can be stimulated to proliferate and exhibit limited neurogenic potential in response to extrinsic cues such as toxic retinal injury, FGF-2 treatment, or modulation of Wnt/β-catenin signaling(12, 15–18). Recent studies have further demonstrated that forced expression of ASCL1 can reprogram Müller glia into neurogenic progenitors capable of generating inner retinal neurons, including bipolar and amacrine-like cells(14, 19). These findings suggest that a latent regenerative program may exist in mammalian Müller cells and can be unlocked under the appropriate molecular conditions. Nevertheless, the neurogenic competence of mammalian Müller cells diminishes with age, largely due to epigenetic constraints. In mature adult mice, chromatin becomes less accessible to pro-regenerative transcription factors. Notably, combining ASCL1 overexpression with epigenetic modulators, such as histone deacetylase inhibitors, can re-establish neurogenic capacity, enabling Müller glia to generate functional neurons that form synaptic connections and exhibit light responsiveness(20).

Translating these advances to humans remains a significant challenge because of species-specific gene expression differences and key anatomical distinctions—most notably the presence of the fovea—that are absent from traditional animal models(21–23). Early investigations using human fetal retina and retinal organoids derived from human pluripotent stem cells have provided proof of concept: human Müller glia can be induced to proliferate and differentiate into neurons exhibiting characteristics of amacrine and ganglion cells following forced ASCL1 overexpression(24, 25). These in vitro models support the notion that human Müller glia is not irreversibly quiescent and may be amenable to reprogramming under appropriate experimental conditions(8).

Using long-term organotypic cultures of adult post-mortem human retinae from 39 donors, we demonstrate that, irrespective of donor age, Müller cell proliferation and the neuronal differentiation of their progeny can be induced by humoral factors(12, 15, 16), without the need for artificial gene expression via viral vectors. Single-cell transcriptomics revealed that cultured Müller glia entered an activated, differentiation-prone state with heterogeneous expression of neurogenic markers. Immunohistochemistry confirmed that postmitotic Müller glia exhibited markers of multiple mature retinal neurons—including photoreceptors, bipolar cells, ganglion cells, amacrine cells, and horizontal cells—demonstrating the intrinsic neurogenic potential of adult human retina.

## Results

### Evidence of Müller cell proliferation in the healthy retina

Physiological cell division in the retina was examined in eyes from organ donors without known ocular disease, enucleated under maintained circulation and promptly fixed to preserve cellular structures. Immunohistochemical labeling of Ki-67, a well-established marker of active proliferation, revealed Ki-67-positive nuclei distributed across multiple layers of the retina **(Fig. S1A)**. To further characterize the proliferating cell populations, additional markers were applied in parallel with Ki-67. Among the Ki-67–positive cells, a substantial proportion were PAX6-positive, indicative of retinal progenitor-like cells, and IBA1-positive, identifying microglia. SOX9 co-expression was observed in only a few Ki-67–positive cells, representing a rare, very limited population of proliferating Müller glia (**Fig. S1B**).

### Induced Müller cell proliferation in long-term organotypic culture of the human retina

To investigate the proliferative and neurogenic capacity of adult human Müller cells, explants from both peripheral and central retinal regions of organ donors with no known ocular disease were maintained in a well-established organotypic culture for up to 10 weeks, in the presence of continuously applied BrdU or EdU nucleoside analogs. During the culture period, the explants maintained the characteristic retinal layering and morphology without showing extensive signs of degeneration.

The in vitro culture environment itself stimulated cell proliferation, with further increases observed at the cut edges of the explants, where the tissue had been exposed to mechanical stress during preparation (**Fig. S2A, B**).

To maximize Müller cell proliferation in culture, we compared treatments with distinct modes of action previously shown to be effective in animal models and continuously administered BrdU or EdU to label proliferating cells. Mild toxicity was mimicked by a single application of 100 µM or 500 µM monosodium glutamate (MSG). The Wnt/β-catenin signaling pathway was activated either by continuous administration of 10 ng/ml or 100 ng/ml recombinant Wnt-3a, or by a single dose of the glycogen synthase kinase-3 inhibitor 6-Bromoindirubin-3′-oxime (BIO) at 1 µM or 5 µM. Finally, cell proliferation was stimulated by continuous administration of 20 ng/ml or 100 ng/ml recombinant FGF-2 (**Fig. S3**). The number of postmitotic cells was quantified in cryosections of large peripheral retinal explants (approximately 1 cm in length) that included the ora serrata. Cell counts were aggregated in 1.5 mm intervals as a function of the distance from the ora serrata and compared to corresponding segments of the same size from central retinal cultures containing the perifoveal region (**Fig. S4; Fig. 1A; Additional file 1)**.

**Fig. 1.**
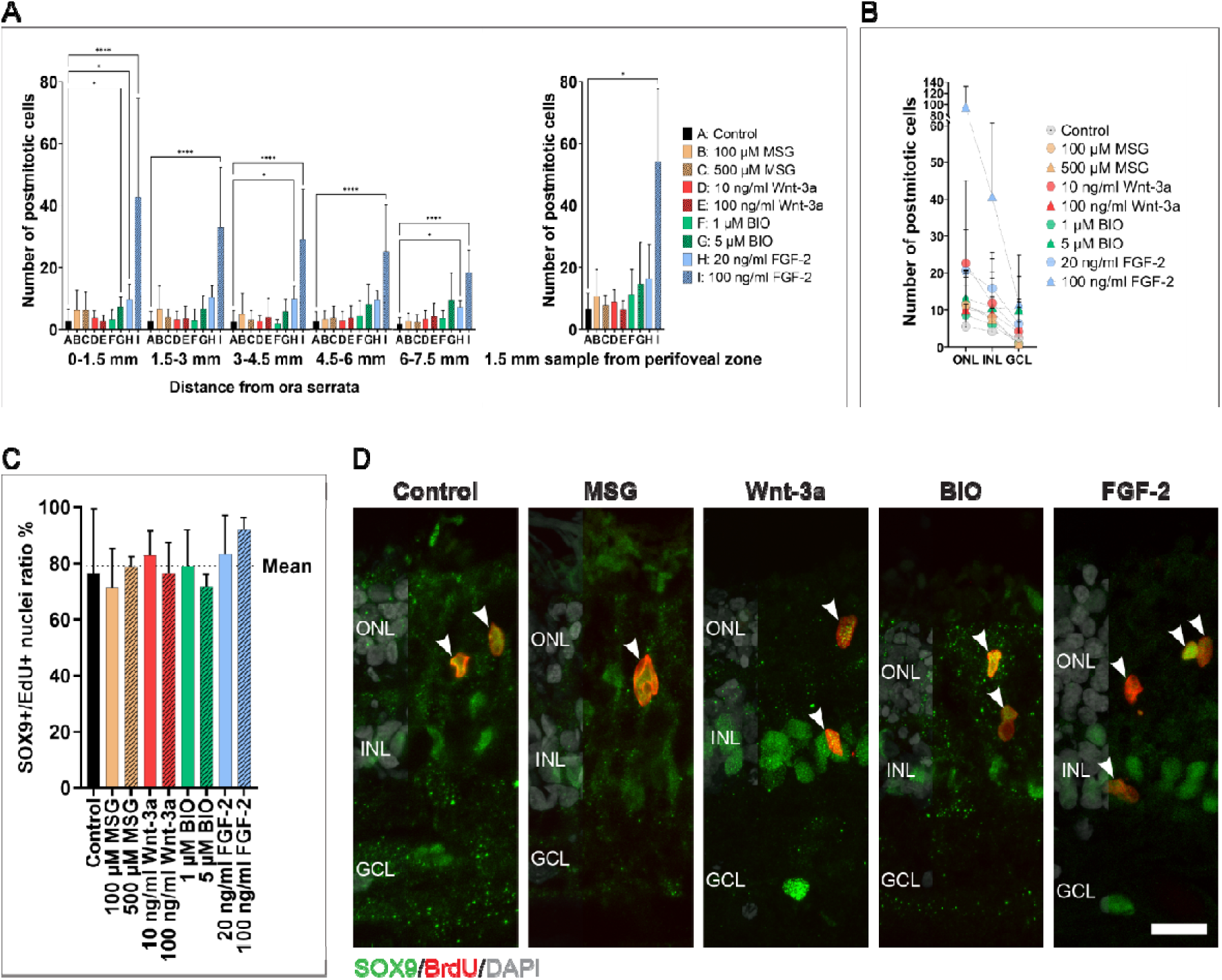
Treatments promote cell division in the adult human retina. **(A)** Postmitotic cell counts in the peripheral (left graph) and perifoveal (right graph) retina. Peripheral measurements were taken in 1.5-mm intervals from the ora serrata toward the center, while the perifoveal region was sampled using a single 1.5-mm segment. Data are shown as mean ± SD. For treatment groups, values from the same donor were averaged before statistical analysis. Shapiro–Wilk testing indicated non-normality in ≥1 group (α = 0.05); therefore, group differences were analyzed by Kruskal–Wallis followed by Dunn’s post hoc test versus control. ****P < 0.0001, ***P < 0.001, **P < 0.01, *P < 0.05. For detailed statistics, see **Additional file 1**. Sample size for peripheral region (left): Control: n = 37 cultures from 20 donors; 100 µM MSG: n = 20 cultures from 8 donors; 500 µM MSG: n = 18 cultures from 8 donors; 10 ng/ml Wnt-3a: n = 13 cultures from 7 donors; 100 ng/ml Wnt-3a: n = 18 cultures from 9 donors; 1 µM BIO: n = 15 cultures from 8 donors; 5 µM BIO: n = 17 cultures from 8 donors; 20 ng/ml FGF-2: n = 8 cultures from 4 donors; 100 ng/ml FGF-2: n = 12 cultures from 6 donors; Sample size for perifoveal region (right): Control: n = 11 cultures from 6 donors; 100 µM MSG: n = 3 cultures from 3 donors; 500 µM MSG: n = 4 cultures from 3 donors; 10 ng/ml Wnt-3a: n = 3 cultures from 3 donors; 100 ng/ml Wnt-3a: n = 3 cultures from 3 donors; 1 µM BIO: n = 3 cultures from 3 donors; 5 µM BIO: n = 3 cultures from 3 donors; 20 ng/ml FGF-2: n = 6 cultures from 3 donors; 100 ng/ml FGF-2: n = 6 cultures from 3 donors. **(B)** Distribution of postmitotic cells across retinal nuclear layers in the periphery. Data are from the same peripheral dataset as Fig. 1A **(left)**. Data are shown as mean ± SD. In the treatment groups, values from the same donor were averaged before plotting. Values were obtained from the peripheral 7.5-mm retinal segment measured from the ora serrata toward the center. For detailed statistics, see **Additional file 1**. **(C)** Proportion of SOX9/EdU co-labeled cells among EdU-positive postmitotic cells in the peripheral retina across control and treated groups. Data are shown as mean ± SD. The combined mean across all treatment groups was 79.09 ± 6.32%, as indicated by the dashed line. Values were obtained from the peripheral 7.5-mm retinal segment measured from the ora serrata toward the center. For detailed statistics, see **Additional file 1**. Sample size: n = 19 cultures from 14 donors; 100 µM MSG: n = 4 cultures from 4 donors; 500 µM MSG: n = 4 cultures from 4 donors; 10 ng/ml Wnt-3a: n = 4 cultures from 4 donors; 100 ng/ml Wnt-3a: n = 4 cultures from 4 donors; 1 µM BIO: n = 4 cultures from 4 donors; 5 µM BIO: n = 6 cultures from 4 donors; 20 ng/ml FGF-2: n = 8 cultures from 4 donors; 100 ng/ml FGF-2: n = 17 cultures from 7 donors. **(D)** Representative images of retinal sections after 10 weeks in culture showing SOX9/BrdU double-positive postmitotic cells in control and treated samples (arrowheads). Sections were immunolabeled for BrdU (red), SOX9 (green), and DAPI (grey). ONL: outer nuclear layer; INL: inner nuclear layer; GCL: ganglion cell layer. Confocal images are shown as z-projections reconstructed from optical sections spanning 3–9 µm, depending on the specimen. Scale bar: 10 μm.

The treatments increased cell proliferation compared with controls, with consistent effects extending from the ora serrata toward the central retina and across the entire retina, irrespective of distance. In peripheral explants, high-dose BIO and both concentrations of FGF-2 significantly increased the level of cell proliferation (**Fig. 1A; Fig. S2B**). In explants from the central retina, a significant effect was observed only with the high-dose FGF-2 treatment (**Fig. 1A; Additional file 1**).

In addition to histological quantification, flow cytometry of Hoechst-positive nuclei revealed an increase in the fraction of EdU-positive cells in peripheral cultures following FGF-2 treatment compared to controls (**Fig. S2C, D; Additional file 1**).

In all conditions examined, the majority of postmitotic cells were located in the outer nuclear layer, followed in frequency by the inner nuclear layer and then the ganglion cell layer (**Fig. 1B; Additional file 1**). 79.09 ± 6.32% of progeny cells were SOX9-positive across all tested conditions, consistent with a Müller glial identity (**Fig. 1C; Additional file 1**). In addition, postmitotic cells showed frequent colocalization with several Müller glia markers, including vimentin, GFAP, S100β, NPVF, and CRYAA (**Fig. 2**).

**Fig. 2.**
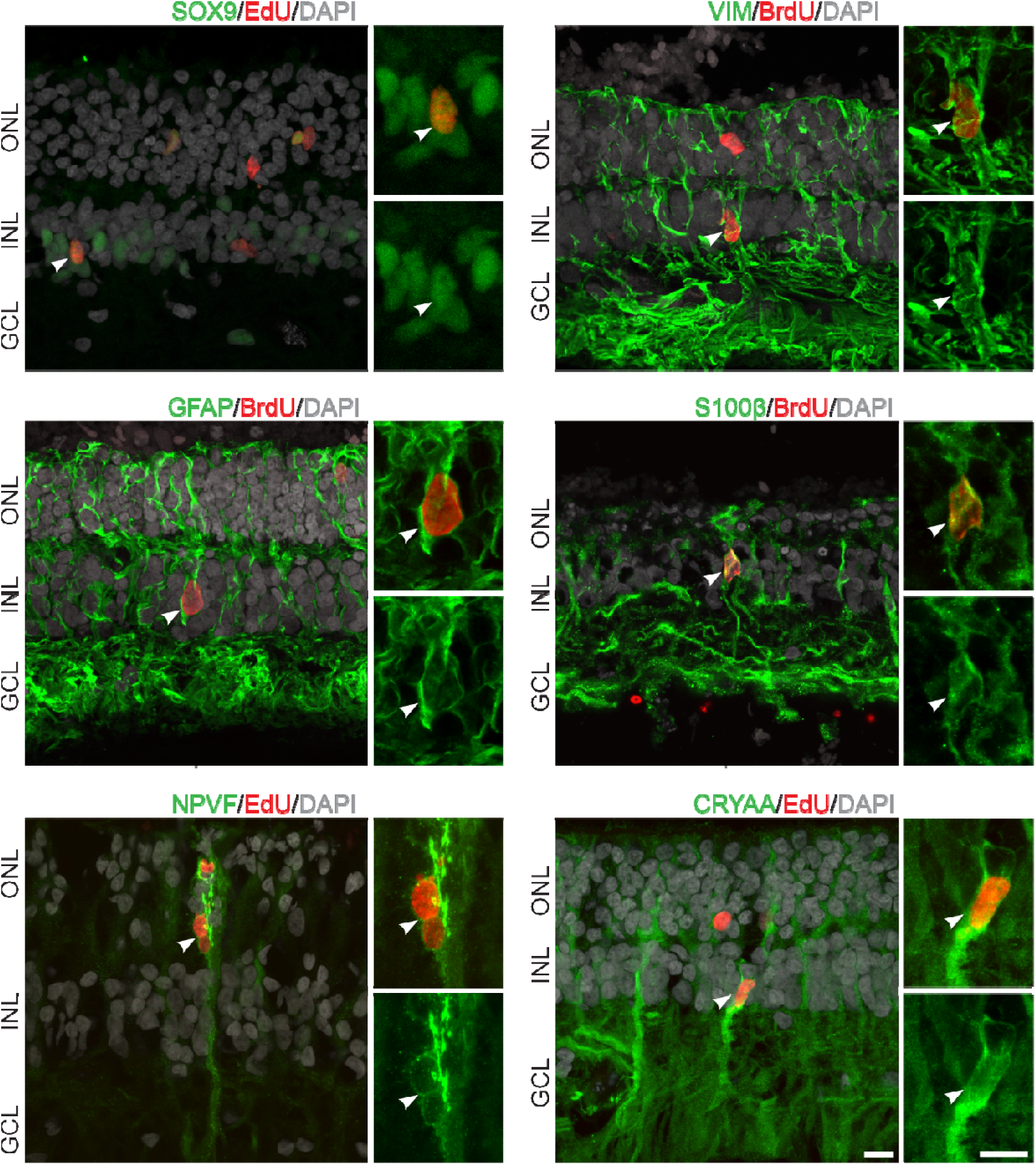
Expression of Müller glia markers in postmitotic cells. Representative images of different induction factor-treated samples showing Müller glia marker expression in postmitotic cells. Immunolabeled 10-week-old retinal sections, white arrowheads indicate the co-labeled cells in both magnifications. Confocal images are shown as z-projections reconstructed from optical sections spanning 1.5–8 µm, depending on the specimen. Scale bars: 10 µm.

The treatments enhanced Müller cell proliferation in both sexes and across all ages, including in older donors, and similar responses were observed across all donors, suggesting that the capacity of human Müller glia to re-enter the cell cycle is a robust and conserved feature of the adult retina. **(Fig. S5; Table S1).**

### Müller glia has the capacity to undergo multiple rounds of proliferation

An EdU/BrdU double-labeling assay was performed to determine whether induced Müller cells can undergo multiple rounds of cell division within their native three-dimensional retinal environment. Alongside FGF-2, high-dose BIO induced the most pronounced increase in proliferation (**Fig. 1A**). Consequently, high concentrations of both compounds were selected for further analysis. We assessed cultures maintained for 2, 5, and 10 weeks, applying both short- and long-term treatment regimens for each factor to analyze cell division dynamics, EdU and BrdU were administered alternately at specific time points to distinguish successive rounds of DNA synthesis (**Fig. S6**). Independent of treatment duration and nucleoside analog switching timing, single-labeled BrdU-positive and EdU-positive cells were consistently observed at all time points across all treatment conditions. At the DIV3 (days in vitro) EdU-BrdU switch, no double-labeled cells were detected. However, beginning with the switch between nucleoside analogs on day 7 and at later time points, small numbers of cells positive for both EdU and BrdU were observed under both BIO and FGF-2 treatment conditions. SOX9 immunolabeling was used to confirm Müller cell identity, and triple colocalization (EdU-positive/BrdU-positive/SOX9-positive) was analyzed. In BIO-treated samples, triple colocalization was detected in a small subset of cells following two consecutive treatment cycles (**Fig. S6A, Conditions #4–#5**). FGF-2 treatment also resulted in triple-positive nuclei positive for all three markers during continuous exposure, as well as cells that had undergone only a single division (**Fig. S6B, Conditions #8–#15; Fig. S6C**). The occurrence of cells that had undergone multiple divisions was not restricted to the ora serrata region or to a particular retinal layer but was observed throughout the retina.

### Long-term culturing and FGF-2 treatment alters Müller cell transcriptional programs

To assess transcriptional changes reflecting Müller glia activation and altered cellular state, single-cell RNA sequencing was performed on untreated retinal cultures and cultures treated with 100 ng/ml FGF-2 for 10 weeks in vitro, using donor-matched samples from three independent donors. These datasets were integrated with a freshly isolated peripheral adult human retina dataset(26) to enable robust, biologically informed cell type annotation using an in vivo reference. Cell types were classified based on the expression of canonical marker genes. Rod photoreceptors were identified by *RHO*, *GNAT1*, and *SAG*, while cones were marked by *ARR3*, *PDE6H*, and *OPN1MW*. Bipolar cells were defined by *GRM6*, *LRTM1*, *TRPM1, PRKCA and GRIK1*. Amacrine cells were characterized by *GAD1* and *SLC6A9*, and horizontal cells were distinguished by *ONECUT1, ONECUT2,* and *LHX1*. Müller cells were identified by markers including *VIM*, *RLBP1*, and *SOX9.* Additional populations included microglia (*CX3CR1*, *AIF1, PTPRC),* astroglia (*GJA1)*, and a small mixed cluster comprising smooth muscle cells (*ACTA2*) and pericytes (*TAGLN*), with potential fibroblasts (*COL1A2*) (**Fig. 3A**). Clusters corresponding to major retinal cell classes were identified in both the cultured and fresh retina datasets in the integrated space (**Fig. 3B**). Rods formed the largest cluster and segregated clearly from cones. Inner retinal neurons resolved into one rod bipolar cluster and six cone bipolar clusters, alongside two amacrine and two horizontal cell clusters. The small mixed cluster with overlapping vascular-associated and fibroblast marker expression was excluded from downstream analysis because it could not be confidently annotated to a single cell type. The Müller glia cluster (including 10723 cells from fresh retina, 4626 from control cultures, and 2970 from treated cultures) was subsetted and re-analyzed independently. Cells were re-integrated and re-clustered prior to differential gene expression analysis (**Fig. 3C**). Re-clustering the Müller glia subset identified six subclusters (**Fig. 3D; Additional file 2**), all of which robustly expressed the canonical Müller glia marker *VIM* (**Fig. 3E**). Genes enriched in these subclusters primarily reflected condition-associated responses—such as metabolic or inflammatory activation and increased protein synthesis—rather than distinct Müller glia subtypes. Therefore, subsequent analyses focused on the total Müller glia population instead of subcluster-level comparisons. Top marker genes are provided in **Additional file 2A**.

**Fig. 3.**
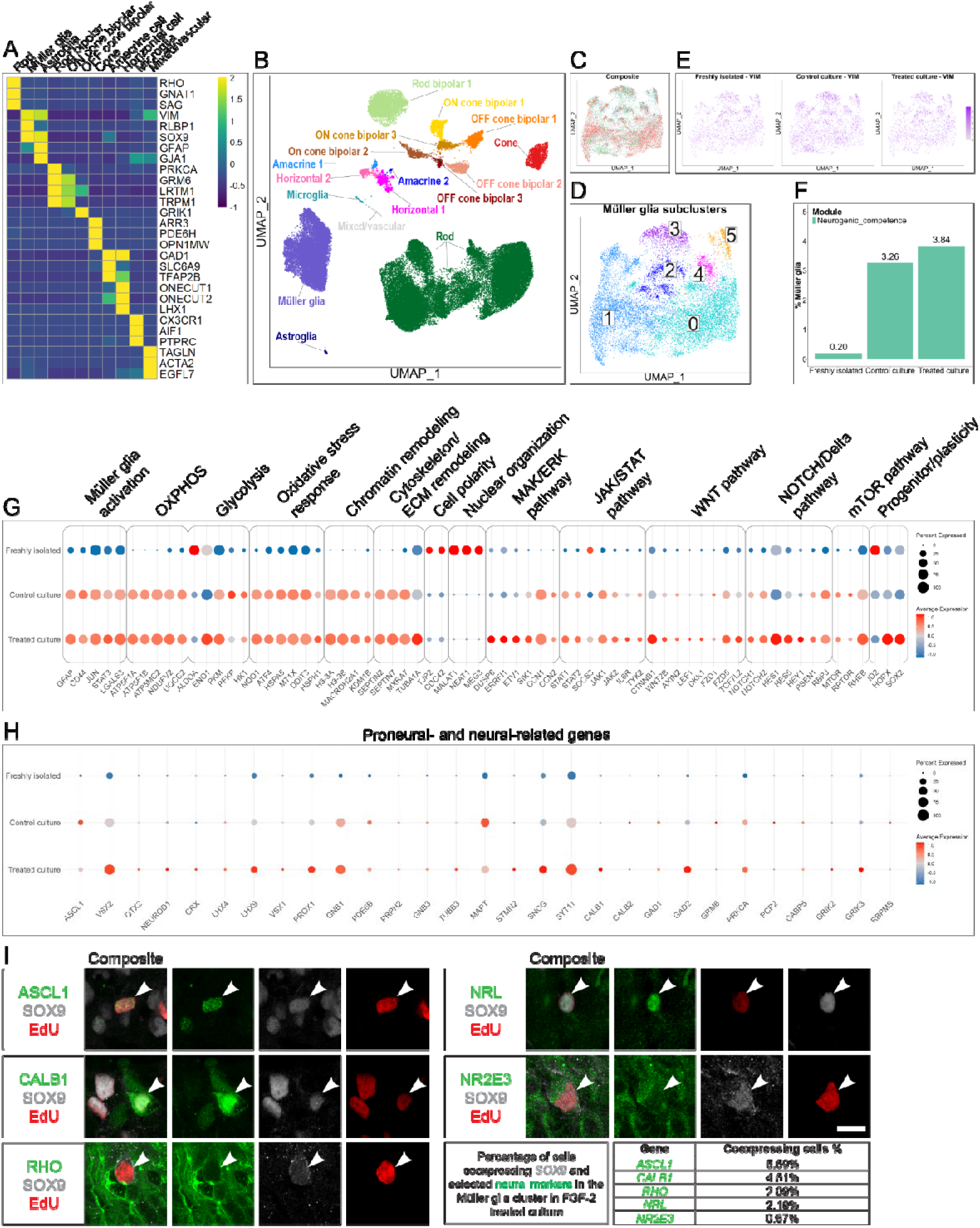
Single-cell RNA-seq reveals culturing and treatment effects in retinal cultures. **(A)** Heatmap of pseudobulk expression profiles showing average normalized expression values of representative marker genes used to identify major retinal and non-retinal cell types in the integrated dataset. **(B)** UMAP visualization of all cells from freshly isolated tissue, control cultures, and FGF-2-treated cultures. Single points represent single cells colored by identity, with major neuronal, glial, and non-retinal populations annotated as identified based on canonical marker gene expression. **(C)** Re-integrated and re-clustered Müller glia subset from freshly isolated retina, control culture, and FGF-2-treated culture groups after harmony integration. (Coral: Freshly isolated, Green: Control culture, Blue: Treated culture) **(D)** Re-integrating and re-clustering identified six Müller glia subclusters. (DEGs in **Additional file 2A - D**) **(E)** Robust *VIM* expression in all subclusters and across experimental groups. **(F)** Bar plot showing the percentage of Müller glia classified with Neurogenic competence across freshly isolated retina, control culture, and treated culture conditions. Neurogenic competence was defined by co-expression of at least three neurogenic/differentiation-associated genes (ASCL1, NEUROD1, NEUROG2, LHX4, LHX9, OTX2, and VSX2) together with Müller glia markers SOX9 and/or VIM. Neurogenic-competent Müller glia were rare in freshly isolated retina (0.20%) but increased in control culture (3.26%) and treated culture (3.84%). **(G)** Dot plot summarizing the transcriptomic differences across pooled Müller glia from all subclusters, across experimental groups. Genes are grouped by functional categories. Dot size represents the proportion of expressing cells, while color represents scaled expression (normalized values centered and scaled per gene). **(H)** Dot plot highlighting transcriptomic patterns associated with reprogramming and neuronal differentiation in Müller glia, showing that cultured cells display transcriptional signatures consistent with neuronal lineage commitment. Dot size represents the proportion of expressing cells, while color represents scaled expression (normalized values centered and scaled per gene). **(I)** Postmitotic SOX9-positive cells co-express proneural- and neural markers in FGF-2-treated human retina. Representative confocal images showing triple-labeled immunofluorescence for the indicated markers (green: ASCL1, CALB1, RHO, NRL, NR2E3), SOX9 (gray), and EdU (red). Immunolabeled 10-week-old retinal sections, white arrowheads indicate the co-labeled cells. Confocal images are shown as z-projections reconstructed from optical sections spanning 1–3.5 µm, depending on the specimen. Scale bar: 10 µm. For more detailed images, see **Fig. S9**.

To quantify overall Müller glia transcriptional similarity between conditions, Jensen–Shannon distances (JSD) were calculated on sample-level pseudobulk expression profiles, a metric ranging from 0 to 1. Control and FGF-2–treated cultures were highly similar (JSD = 0.116), while both diverged substantially from Müller glia derived from freshly isolated retina (JSD = 0.419, 0.412, respectively; **Fig. S7**). Culturing induced pronounced transcriptional changes across multiple biological processes, with additional, subtle but specific modulation by FGF-2 treatment (**top between-group DEGs reported in supplementary Additional file 2B, C**). Cultured Müller cells showed elevated expression of markers associated with reactive gliosis, including GFAP, CD44, and *LGALS3,* together with activation*-*associated transcription factors such as *JUN* and *STAT3*. Genes involved in metabolic activity and oxidative stress responses were strongly upregulated in cultured conditions (**Fig. 3G; Additional file 2B, C**), for example, *NQO1* (log2FC = 4.73, p.adj = 5.92×10^-205^; reported statistics reflect control culture vs. freshly isolated retina unless stated otherwise). Similarly, genes associated with cytoskeletal remodeling (e.g. *SEPTIN7*: log2FC = 7.14, p.adj = 9.31×10^-155^), chromatin regulation (e.g. *H3-3A*: log2FC = 8.21, p.adj = 8.95×10^-168^, and stress responses (e.g. *CCN1*: log2FC = 8.41, p.adj = 3.88×10^-176^) were similarly upregulated in cultured conditions, with FGF-2 treatment further enhancing differentiation and neuronal fate potential (such as *ETV1, DUSP6, CALB1,* and *GAD2*) (**Fig. 3G**; **Additional file 2D**). In contrast, a subset of genes involved in establishing cell polarity and nuclear organization were significantly downregulated in culture compared to freshly isolated retina, including *TJP2* (log2FC = −2.61, p = 4.18×10^-105^) and *NEAT1* (log2FC = −6.08, p = 3.63×10^-78^. These alterations may reflect, at least in part, differences between the in vivo retinal milieu and ex vivo culture conditions. **Fig. 3G; Additional file 2B**).

More subtle transcriptional changes were also observed in cultured Müller glia in components of progenitor identity and differentiation-related signaling pathways, including genes involved in JAK/STAT, WNT, NOTCH/Delta, and mTOR signaling pathways. Expression of progenitor maintenance-associated regulators, such as *SOCS3* and *ID2*, was expressed at lower levels in culture (log2FC = −1.98, p.adj = 3.32×10-05; log2FC = −2.86, p.adj = 1.27×10^-17^, respectively, **Fig. 3G**). These patterns suggest that the culture environment favors an activated, differentiation-permissive state rather than progenitor maintenance(27–29). Together, these transcriptional shifts indicate that ex vivo culturing induces a pronounced reactive, metabolically active, and differentiation-prone state in Müller glia, with FGF-2 treatment selectively amplifying components of this response. These observations are consistent with our immunohistochemical findings of increased expression of progenitor-associated markers, such as PAX6 and SOX2, in treated cultures (**Fig. S8**).

### Induction of Müller glia results in neural marker expression in their progeny

To evaluate whether metabolic activation in cultured Müller glia was accompanied by neurogenic reprogramming, we examined the expression of early proneural transcription factors and lineage-associated markers, with selected targets validated at the protein level (**Fig. 3H, I; Fig. S9; Table S2**). Notably, early neurogenic progenitor regulator *ASCL1* was elevated even in control conditions in culture compared to fresh retina (log2FC = 2.21, p.adj = 0.0037), suggesting baseline activation of neurogenic competence in culture conditions. Robust *ASCL1* expression was also confirmed on the protein level in cultured, post-mitotic, SOX9-positive cells (**Fig. 3I; Fig. S9**). In contrast, bipolar lineage regulator *VSX2* was significantly upregulated only in FGF-2-treated cultures (log2FC = 0.93, p.adj = 0.0012), suggesting a modest enhancing effect of the treatment on neurogenic activation. Additional progenitor-associated factors, including *OTX2* and *NEUROD1,* were detected at low frequencies in both cultured conditions but did not reach statistical significance. To enable more robust comparison of neurogenic potential across groups, we defined a neurogenic, differentiation-associated gene module (including *ASCL1*, *NEUROD1*, *NEUROG2*, *LHX4*, *LHX9*, *OTX2*, and *VSX2*) and quantified the percentage of Müller glia co-expressing at least three of these genes together with the Müller markers *SOX9* and/or *VIM*. In line with our observations, such cells represented 3.26% of Müller glia in control cultures, 3.84% in treated cultures, but were detected only rarely in Müller glia clusters from freshly isolated retina (0.20%, **Fig. 3F**).

Genes associated with intermediate stages of retinal cell fate specification (such as *CRX, LHX4*, and LHX9), as well as late-stage fate regulators (such as *VSX1* and *PROX1*), were also detected, suggesting the activation of lineage commitment–associated programs within cultured Müller cells. Consistent with this observation, genes associated with neuronal maturation and function—including *TUBB3*, *CALB1*, and *CALB2*—were also observed at heterogeneous levels, indicating activation of neuronal transcriptional programs within cultured Müller glia. Markers associated with mature retinal neuronal cell types were sporadically observed, including bipolar cell genes such as *GRM6*, *PRKCA*, *PCP2* and *CABP5*, amacrine cell markers such as *GAD1* and *GAD2,* and ganglion cell marker *RBPMS* (**Fig. 3H**; **Table 1; Table S2**). Notably, *GAD2* expression was significantly higher in FGF-2-treated cultures compared to freshly isolated retina (log2FC = 2.89, p.adj = 0.012) and even compared to control cultures (log2FC = 1.18, p.adj = 2.26×10^-06^), potentially suggesting enhanced activation of amacrine-like transcriptional programs under growth factor stimulation (**Fig. 3H**).

**Table 1.** Percentage of cells in the total Müller glia cluster expressing genes depicted in morphowlogical images (**Fig. 4**, **Fig. 5)**

| Percentage of cells in the total Müller glia cluster expressing: |  |  |  |
| --- | --- | --- | --- |
| Gene | Freshly isolated | Control culture | Treated culture |
| *PAX6* | 63.723% | 72.698% | 81.246% |
| *SOX2* | 66.605% | 67.401% | 73.737% |
| *ASCL1* | 1.301% | 14.786% | 8.653% |
| *LHX4* | 0.000% | 0.994% | 1.448% |
| *VSX2* | 25.170% | 56.572% | 71.582% |
| *TUBB3* | 0.065% | 0.735% | 1.313% |
| *ELAVL3* | 0.187% | 0.367% | 0.606% |
| *ELAVL4* | 0.187% | 0.000% | 0.101% |
| *SCGN* | 0.037% | 0.000% | 0.034% |
| *CALB1* | 1.222% | 0.043% | 5.623% |
| *CALB2* | 0.224% | 0.259% | 0.134% |
| *CRX* | 0.625% | 0.238% | 0.606% |
| *NR2E3* | 0.214% | 1.513% | 0.943% |
| *NRL* | 7.078% | 1.729% | 3.602% |
| *GNAT1* | 8.608% | 2.789% | 5.319% |
| *RHO* | 9.186% | 9.338% | 3.333% |
| *ARR3* | 1.100% | 0.043% | 0.000% |
| *OPN1SW* | 0.177% | 0.195% | 0.303% |
| *OPN1LW* | 0.625% | 0.000% | 0.000% |
| *OTX2* | 0.103% | 0.303% | 0.438% |
| *PRKCA* | 7.927% | 15.585% | 17.070% |
| *RBPMS* | 0.046% | 0.346% | 0.337% |
| *BRN3A* | 0.000% | 0.000% | 0.000% |
| *RBFOX3* | 0.046% | 0.518% | 0.269% |
| *SYP* | 0.000% | 0.367% | 1.751% |

Photoreceptor-expressing cells were also present; however, interpretation is confounded by potential residual ambient RNA contributions. Nevertheless, protein-level co-detection of SOX9 and EdU together with rod photoreceptor markers RHO, NRL, and NR2E3 supports the existence of rare proliferative Müller glia–derived cells exhibiting rod photoreceptor–like features (**Fig. 3I**; **Fig. S9**; **Table 1; Table S2**). Throughout the analysis, multiple rounds of rigorous filtering were applied. Cells within the Müller glia subpopulation that co-expressed the observed neural markers retained at least a partial Müller glia identity and consistently expressed canonical markers such as *VIM* and *RLBP1*, confirming that the observed transcriptional changes arose within the Müller glia lineage.

### Immunohistochemical validation of neuronal commitment in Müller glia-derived progeny

Immunohistochemistry was performed on retinal cryosections to confirm the observed transcriptomic patterns at the protein level. To assess whether proliferating cells initiated neural differentiation, colocalization analyses were carried out using a panel of progenitor- and proneural-specific markers (**Fig. 4**; **Table 1**). FGF-2 treatment led to marked upregulation of PAX6 and SOX2, with a large fraction of postmitotic cells co-expressing these markers (**Fig. 4, Fig. S8**). In contrast to the fresh retina, ASCL1 expression was observed in a large subset of EdU-positive cells in both control and FGF-2–treated cultures. Numerous EdU-positive postmitotic cells expressed LHX4 and VSX2 (also known as CHX10), which are well-known markers associated with retinal progenitors and neuronal differentiation (**Fig. 4**).

**Fig. 4.**
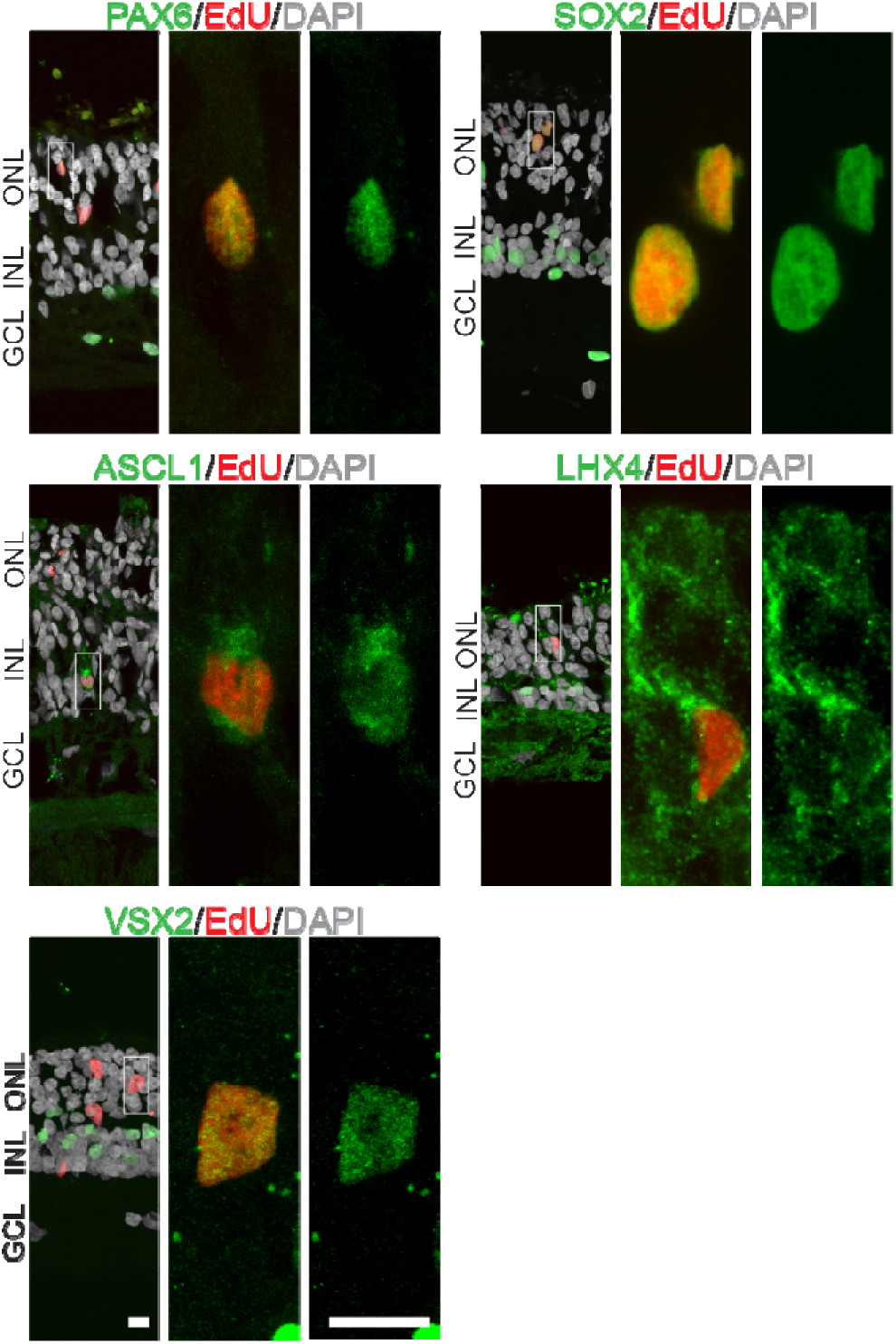
Expression of progenitor and proneural markers in postmitotic cells. Representative images of high-dose FGF-2–treated retinal sections (10-week-old cultures) showing expression of progenitor and proneural markers (PAX6, SOX2, ASCL1, LHX4, VSX2) in EdU-positive postmitotic cells. Left panels show overview images, with white boxes indicating the position of the cell shown at higher magnification in the corresponding middle and right panels. Confocal images are shown as z-projections reconstructed from optical sections spanning 1.5–5.5 µm, depending on the specimen. Scale bars: 10 µm. Percentage of cells in the total Müller glia cluster expressing the depicted markers in **Table 1**.

To determine whether these cells progressed toward neuronal maturation, we next examined markers of differentiated retinal neurons (**Fig. 5**; **Table 1**). A large proportion of postmitotic EdU-positive cells expressed neuronal cytoskeletal marker β-tubulin III (TUBB3) and the HuC and HuD proteins (encoded by the *ELAVL3* and *ELAVL4* genes, respectively), as a sign of neuronal commitment. A small population of EdU-positive cells expressed secretagogin (SCGN), indicating developing or mature amacrine or bipolar cells. Markers associated with additional retinal neuronal subtypes were also detected among newly generated postmitotic cells. These included calretinin (CALB2) and calbindin (CALB1), proteins known to be expressed in subsets of horizontal, amacrine, and ganglion cells in the human retina. A small number of EdU-positive cells expressed photoreceptor-specific proteins, including CRX and recoverin, as well as rod lineage markers NR2E3, NRL, transducin alpha-1 chain (GNAT1), and rhodopsin (RHO). Cone-associated markers, including arrestin-3 (ARR3), S-opsin (OPN1SW), and M/L-opsin (OPN1MW/ OPN1LW), which are typical of mature photoreceptor subtypes, were also detected. Additionally, expression of OTX2, PKCα revealed commitment toward bipolar cell differentiation. Postmitotic EdU-positive cells with potential ganglion cell identity were also identified by immunolabeling for RBPMS, BRN3A, and NeuN. Finally, the presynaptic protein synaptophysin was detected in multiple postmitotic cells, with representative examples showing co-expression with neuronal markers such as calbindin and arrestin-3, indicating the acquisition of synaptic protein expression in a subset of newly generated cells (**Fig. S10**).

**Fig. 5.**
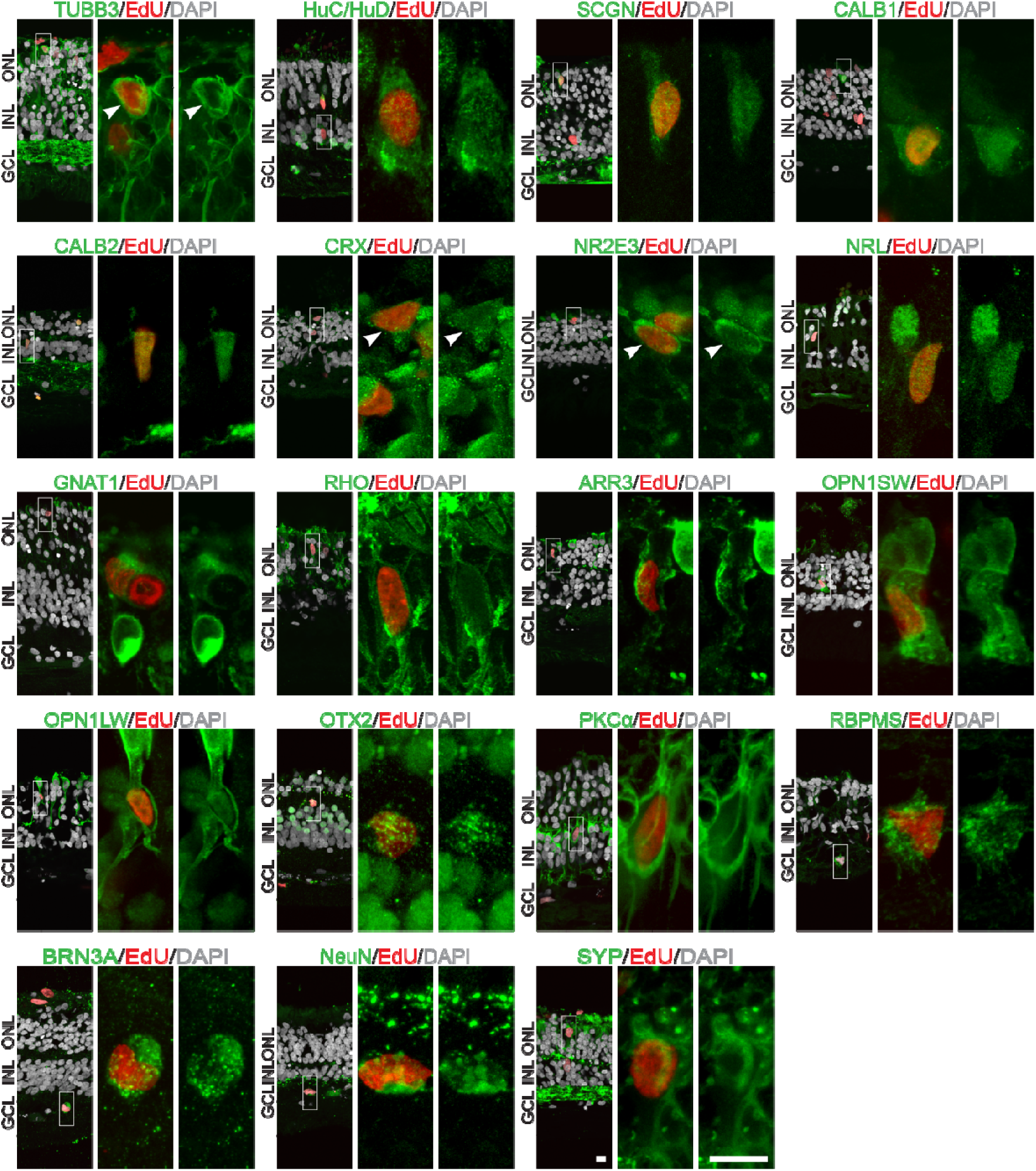
Expression of neural markers in postmitotic cells. Representative images of high-dose FGF-2-treated retinal sections (10-week-old cultures) showing expression of neural markers in EdU-positive cells. The left panels show overview images, with white boxes indicating the position of the cell shown at higher magnification in the corresponding middle and right panels. Arrowheads indicate colocalized cells; multiple EdU-positive cells are displayed. Confocal images are shown as z-projections reconstructed from optical sections spanning 1.5–5 µm, depending on the specimen. Scale bars: 10 µm. Percentage of cells in the total Müller glia cluster expressing the depicted markers in **Table 1**.

## Discussion

Recent advances have broadened the repertoire of human retinal tissue models(30–32), yet important limitations persist. Retinal organoids and fetal retina–derived cultures enable long-term studies, but they do not fully recreate the cellular diversity, maturation state, or physiological environment of the adult retina. Consequently, data obtained from these systems must be interpreted with an understanding of their developmental bias. Post-mortem retinal tissue from adult donors remains the most faithful model for studying adult retinal biology, although its utility is constrained by limited access to high-quality tissue and the technical challenges of sustaining adult explants over extended periods. Here, we successfully implemented our proven post-mortem human adult retina culture system, which supports complex, long-term experimentation in adult human retinal tissue, thereby helping to bridge this gap and expanding the experimental possibilities for adult-relevant retinal research(33–35).

Retinas collected from ophthalmologically healthy organ donors under maintained circulation and fixed immediately contained SOX9-positive dividing cells, pointing to a residual proliferative capacity in Müller glia that persists in the adult human retina under physiological conditions. Long-term cultures established from these donor retinas showed a pronounced rise in the number of Müller cells progressing into the cell cycle. The increase in proliferative activity was most prominent at the cut margins of the cultures, consistent with the significant mechanical stress these regions sustained during explant preparation. Across samples from central and peripheral retinal areas, the distribution of proliferating cells was remarkably consistent. These findings suggest that inducible Müller cells are present throughout the adult human retina, rather than being limited to the extreme periphery or to areas beyond the ora serrata, where progenitor cells reside in the ciliary marginal zone in certain species(36). The humoral factors applied to induce proliferation further increased the number of cell divisions in the treated cultures. Among these factors used to stimulate proliferation, high doses of the GSK-3 inhibitor BIO and FGF-2 were especially effective, leading to an additional increase in proliferative activity within the treated cultures prepared from both peripheral and central retinal regions. Examination of several hundred explants derived from 39 donors aged 18 to 68 years demonstrated that Müller cell proliferation is not limited to young individuals; rather, it can be induced across the adult age spectrum in both sexes.

We observed a higher prevalence of postmitotic cells in the ONL compared to other layers at the end of the culturing period. This phenomenon may be reminiscent of interkinetic nuclear migration described in lower vertebrates(37, 38). However, the dynamics of nuclear movement cannot be determined from our experiments. The accumulation of postmitotic cells within the ONL likely reflects the culmination of proliferative events followed by differentiation processes that remain to be fully characterized in the adult human context. Notably, the spatial distribution of these cells did not correspond to any obvious gradients in tissue viability or treatment responsiveness, suggesting that their localization is driven by intrinsic properties of Müller glia–derived progenitors rather than by extrinsic culture artifacts.

The consistently high (79.09 ± 6.32%) proportion of SOX9-positive progeny cells, as well as their co-expression with several Müller glia markers across all examined conditions, supports the conclusion that Müller cells constitute both the primary source of proliferating cells and the cell type produced by proliferation. This finding further suggests that the majority of postmitotic cells retain Müller cell identity and exhibit only limited differentiation potential under the experimental conditions used.

Using an EdU/BrdU double-labeling assay, we found that most postmitotic cells incorporated only a single nucleoside analog, suggesting either that only one division occurred during the culture period or that additional divisions occurred within a narrow time window not captured by the switch between the two labels. However, we also confirmed that Müller cells are, in principle, capable of undergoing multiple rounds of proliferation following induction, as evidenced by a subset of cells that incorporated both EdU and BrdU. This indicates that while the overall proliferative response is limited for the majority of Müller cells, a fraction of the population can sustain repeated cell-cycle re-entry under the applied experimental conditions.

In the scRNA-seq dataset, Müller glia from freshly isolated human retina displayed low expression of reactive gliosis markers and metabolic stress genes, consistent with their quiescent in vivo state in the healthy retina. This was altered in culture, where in vitro control conditions alone induced a reactive Müller glia phenotype, characterized by upregulation of gliotic and stress-response genes. These changes indicate that culturing triggers a metabolically demanding, stress-associated state in Müller cells. Beyond alterations in gliosis-related gene expression, the culture conditions also led Müller glia to upregulate genes associated with neural commitment and differentiation. FGF-2 treatment further amplified this effect, promoting neural lineage commitment in a subset of progeny cells. At the gene-expression level, we observed modulation of WNT and mTOR pathway components, sustained JAK/STAT activation, and partial suppression of NOTCH/Delta signaling in Müller glia clusters. These molecular changes are consistent with observations from mammalian and human tissue models, as well as with findings showing that FGF signaling promotes Müller glia dedifferentiation and neural regeneration(27, 39, 40).

We further investigated whether Müller cell–derived progeny acquire neural characteristics and found that a subset of proliferating cells expressed progenitor and proneural markers, including ASCL1, a key factor in Müller glia reprogramming identified in mammalian studies(20, 27). The majority of newly generated progeny remained SOX9-positive; however, a subset of these cells also expressed different neural markers, indicating that neural commitment can begin while nuclear SOX9 expression is still maintained. We next examined whether this postmitotic population expressed markers associated with terminal neuronal differentiation. Previous studies of Müller cell reprogramming reported that progeny marker expression was restricted to a limited subset of neuronal cell types(25). In contrast, the analyzed Müller glia clusters, which contained a large proportion of the progeny cells examined here, showed expression of markers corresponding to all major retinal neuronal classes at both the mRNA and protein levels, with spatial patterns consistent with the physiological locations of the respective mature cell types. The immunohistochemically verified expression of several late-stage photoreceptor proteins—including CRX, recoverin, NR2E3, arrestin-3, GNAT1, rhodopsin, S-opsin, and M/L-opsin—clearly indicates that, in a subset of progeny cells located in the ONL, the development of both cones and rods took place. In addition, within the inner nuclear layer, a subset of postmitotic cells exhibited expression of interneuron-associated markers, including OTX2, PCP2, PKCα, calretinin, and calbindin, while in the ganglion cell layer, a portion of the BrdU- or EdU-positive cells expressed markers characteristic of mature ganglion cells, such as RBPMS, NeuN, and BRN3A. Although the functionality of the newly generated neurons and their proper integration into the retinal network could only be confirmed definitively by electrophysiological analyses, the co-presence of synaptophysin in cells expressing markers characteristic of mature neurons suggests that early processes required for the initiation of synapse formation may be underway.

Our findings highlight that Müller glia in the adult human retina—regardless of age or sex—are capable of re-entering the cell cycle, undergoing multiple rounds of division, and committing to all major neural cell lineages even in the absence of targeted ASCL1 overexpression via viral vectors. Future studies may enable the optimization of conditions for Müller glia–derived neurogenesis and the investigation of the functional integration of newly generated cells, which could ultimately support the development of regenerative therapies for the treatment of degenerative retinal diseases.

## Materials and Methods

Note: Reference numbers for reagents are listed in **Table S3.**

### Human retinal tissue

78 eyes from 39 donors, aged 18-68 years, without known ophthalmic disease, were obtained via multi-organ donation (**Table S4**). All eyes were enucleated prior to cardiac arrest in accordance with the Declaration of Helsinki. All experimental protocols were approved by the National Ethics Committee (ETT TUKEB 34851-2/2018/EKU and ETT TUKEB IV/5645-1/2021/EKU).

### Human retinal cultures

Following enucleation, the cornea was immediately removed and preserved for transplantation, while the remaining eyecup was transported to the laboratory in chilled DMEM/F-12 medium. In the laboratory, the eyes were dissected under a stereomicroscope, and the neural retina was isolated. Subsequently, 4×4 mm central or peripheral pieces of neural retina and 5×10 mm centro-peripherally oriented segments containing the ora serrata were isolated and placed on polycarbonate membrane culture inserts (**Fig. S4**). The cultures were maintained in DMEM/F-12 medium (Gibco) supplemented with 0.1% BSA (Millipore), 10 μM O-acetyl-l-carnitine hydrochloride, 1 mM fumaric acid, 0.5 mM galactose, 1 mM glucose (Merck), 0.5 mM glycine, 10 mM HEPES, 0.05 mM mannose, 13 mM sodium bicarbonate, 3 mM taurine, 0.1 mM putrescine dihydrochloride, 0.35 μM retinol, 0.3 μM retinyl acetate, 0.2 μM (±)-α-tocopherol, 0.5 mM L-ascorbic acid, 0.05 μM sodium selenite, 0.02 μM hydrocortisone, 0.02 μM progesterone, 1 μM insulin, 0.003 μM 3,3′,5′-triiodo-l-thyronine, 2000 U penicillin and 2 mg streptomycin at 37°C with 5% CO₂. The medium was changed every two days, and cultures were maintained for up to 70 days.

Proliferating cells were tracked using 5 mg/ml 5-bromo-2’-deoxyuridine (BrdU) and 10 µM 5-ethynyl-2’-deoxyuridine (EdU), which were continuously added to the culture medium. To stimulate cell division, various humoral factors with different mechanisms of action were added to the culture medium under different conditions. Monosodium glutamate (MSG) was applied at concentrations of 100 mM and 500 mM on DIV1 (days in vitro) for 60 minutes (min), after which the medium was replaced. Recombinant human Wnt-3a (R&D systems) was added at concentrations of 10 ng/ml and 100 ng/ml to the cultures on DIV1 for 48 h, followed by a medium change. The glycogen synthase kinase-3 (GSK-3) inhibitor (2’Z,3’E)-6-Bromoindirubin-3’-oxime (BIO) (Tocris) was applied at concentrations of 1 μM and 5 μM on DIV1 for 48 hours, followed by a medium change. Human recombinant Fibroblast growth factor 2 (FGF-2) was continuously added to the medium at concentrations of 20 ng/ml and 100 ng/ml. All chemicals were obtained from Sigma-Aldrich unless otherwise stated (**Table S3**).

### Immunohistochemical analysis

#### Tissue preparation for immunofluorescence staining

Fresh human eyecups used for the assessment of mitotic activity in vivo were fixed immediately after enucleation and cornea removal in 4% paraformaldehyde (PFA) prepared in 0.1 M phosphate buffer (PB, pH 7.4) for 2 h at room temperature (RT). Cultures were fixed under identical conditions for 30 min. Following fixation, the samples were washed three times in 0.1 M PB for 10 min each, transferred to 30% sucrose prepared in 0.1 M PB, and incubated overnight at 4°C. The samples were subsequently frozen and stored at −80°C until further use.

#### Whole-mount immunohistochemistry of retinal explants

Fixed human retinal explants, including both freshly fixed and cultured tissues, were subjected to three freeze-thaw cycles in 30% sucrose prepared in 0.1 M PB on dry ice. Samples were then washed for 15 min in 0.05 M phosphate-buffered saline (PBS, pH 7.4) and blocked overnight at RT in blocking solution (BS; 1% bovine serum albumin (BSA), 0.4% Triton X-100, and 0.05% sodium azide in PBS). Primary antibodies (**Table S3**) were diluted in BS and applied for 3 days at RT on a rocking platform. Tissues were subsequently washed three times in PBS and incubated overnight at RT with secondary antibodies and 4′,6-diamidino-2-phenylindole reagent (DAPI; Thermo Fisher Scientific) diluted in BS. EdU incorporation was visualized by click chemistry according to the manufacturer’s protocol (BaseClick). Samples were mounted on Superfrost Plus slides (Thermo Fisher Scientific) in 90% glycerol/PBS, with the ganglion cell layer oriented toward the coverslip.

#### Immunohistochemistry on cryosections

Cultured human retinal explants were embedded in OCT medium, cryosectioned at 20–30 μm, mounted on gelatin- or silane-coated slides, and dried overnight at 37°C. Sections were used immediately or stored at –20°C. Following rehydration in PBS, samples were incubated in BS for 2 h at RT. Primary antibodies were diluted to specific concentrations in BS (**Table S3**) and applied overnight at 4°C. The slides were then washed three times with PBS and incubated with secondary antibodies diluted in BS for 2 h at RT. BrdU detection was performed prior to immunohistochemistry. For this, slides were treated with 2.5 µg/ml Proteinase K solution at RT for 5 min and washed twice with Phosphate-Buffered Saline + Tween-20 (PBST, 0,1% Tween-20 in PBS) containing 2 mg/ml glycine. This was followed by two additional washes with PBST, after which samples were post-fixed in 4% PFA for 20 min at RT. The samples were then washed three times with pre-warmed DNase buffer (21 mM MgCl₂ in 50 mM Tris(hydroxymethyl)aminomethane hydrochloride (TRIS-HCl), pH 7.5) and incubated with DNase solution (0.25 mg/ml DNase I in DNase buffer) at 37°C. Subsequently, they were fixed in pre-chilled 4% PFA on ice for 20 min, incubated in cold TRIS-HCl (50 mM, pH 7.5) buffer for 5 min, and washed three times with PBS. The anti-BrdU antibody was applied according to the standard immunohistochemistry protocol. EdU incorporation was visualized using a click reaction (BaseClick), performed according to the manufacturer’s instructions. Finally, slides were mounted with Fluoroshield mounting medium containing DAPI.

### Microscopy and quantification of cell count

Confocal z-stacks were acquired using a Zeiss LSM 780 confocal microscope equipped with ZEN software (ZEN 2012, Zeiss) and 10×, 20×, 40×, or 63× objectives. Confocal z-stacks were merged using ImageJ (Fiji distribution), and further linear image processing was performed in Photoshop (Adobe). BrdU- and EdU-positive cells were quantified manually from processed images. Donor datasets used for cell counting were pooled to increase the robustness of the analysis. Detailed acquisition and processing parameters are provided in the figure legends.

### Assessing age- and gender-dependent variation in mitotic induction

To compare the effects of age and sex on treatment-induced cell division, a mitotic index was defined for each donor. Under high-dose treatment conditions, the mean number of nucleoside analog–positive cells per unit volume was calculated for each donor. To allow comparison of the degree of mitotic induction between donors, this donor-specific mean was normalized by dividing it by the mean number of nucleoside analog–positive cells per equivalent volume in untreated control samples pooled from all donors within the same treatment group.

### Dissociation and flow cytometry

#### Dissociation and fixation

Retinal tissues were dissociated using the Neural Tissue Dissociation Kit (P) (Miltenyi Biotec). Mid-peripheral 4×4 mm pieces of 10-week-old cultured human retina were placed in a 1.5 mL tube containing PBS. Enzyme P mix was prepared by combining 40 μl of Enzyme P with 950 μl of Buffer X. The PBS was removed, and the Enzyme P mix was added to the tissue. Samples were incubated at 37°C on a temperature-controlled shaker at 900 rpm for 30 min. After the initial incubation, 10 μl of Enzyme P was added, followed by a further 10 min incubation under the same conditions. Next, 15 μl of Enzyme A mix (5 μl Enzyme A + 10 μl Buffer Y) was added, and the samples were incubated for an additional 10–15 min at 37°C. Following incubation, the tubes were placed on ice for 1-2 min. Samples were then centrifuged at 300 × g for 5 min at 4°C. The supernatant was removed, and the pellet was resuspended in 500 μl of staining buffer (BD Biosciences, supplemented with 5 mM EDTA). The suspension was filtered sequentially through a 70 μm filter and then a 40 μm filter. The filtrate was centrifuged at 400 × g for 5 min at 4°C. After centrifugation, 400 μl of supernatant was discarded, leaving a 100 μl cell suspension for fixation. Cells were fixed and permeabilized by adding 100 μl of Foxp3 Fixation/Permeabilization working solution (eBioscience Intracellular Fixation & Permeabilization Buffer Set), prepared according to the manufacturer’s instructions. The mixture was pulse-vortexed and incubated for 30 min at RT, protected from light. Following incubation, 1 mL of 1× Permeabilization Buffer was added, and samples were centrifuged at 400 × g for 5 min at RT. The supernatant was discarded, and the pellet was resuspended in the residual 100 μl volume.

#### EdU detection

The EdU click reaction cocktail was prepared freshly according to the cryosection EdU detection protocol (100 µl/tube). The EdU reaction mix was added to the samples, and the suspension was incubated for 30 min at RT, protected from light. Subsequently, the tubes were filled with 1X Permeabilization Buffer, and the samples were centrifuged at 500 × g for 5 min at RT. The supernatant was discarded, and the pellet was resuspended in 100 µl of the residual volume. This washing step was repeated one more time. Hoechst (diluted 1:250) was added to the residual suspension and incubated for 30 min at RT under light-protected conditions for nuclear staining. The tubes were filled with 1X Permeabilization Buffer, and the samples were centrifuged at 500 × g for 5 min at RT. The supernatant was discarded, and the pellet was resuspended in 100 µl of the residual volume. This washing step was repeated one more time. Finally, the samples were resuspended in 200 µL of staining buffer for analysis. Flow cytometric acquisition was performed using a CytoFLEX (Beckman Coulter). At least 20000 events were acquired per sample when possible. In cases of limited cell yield, all available events were collected and included in the analysis. Data was processed using FlowJo software (Tree Star, version 10.10).

### Single-Cell RNA Sequencing

#### Single-Cell RNA-Seq Library Preparation

Retinal explants cultured for 10 weeks were dissociated into single-cell suspensions following the same protocol employed for flow cytometry experiments. A total of 8000 dissociated cells were loaded onto a 10x Genomics Chromium Next GEM Chip G. Single-cell RNA sequencing (scRNA-seq) libraries were prepared using the Chromium Next GEM Single Cell 3′ Reagent Kits (version 3.1) in accordance with the manufacturer’s instructions (manual version CG000204_Rev_C). Indexed sequencing libraries were subsequently sequenced on an Illumina HiSeq 2500 platform. For comparative analysis, our cultured scRNA-seq data were integrated with a previously published human retinal dataset(26).

#### Data preprocessing and quality control

Raw BCL files from cultured samples were demultiplexed using Cell Ranger *mkfastq* (v2.0/v3.1), and reads were aligned to the human reference genome (GRCh38) using *count* to generate raw and filtered gene–cell count matrices. An existing scRNA dataset from adult human freshly isolated light-sensitive retina was obtained from the European Genome-phenome Archive (EGAS00001004561)(26). BAM files were converted to FASTQ using Cell Ranger *bamtofastq* and preprocessed using the same pipeline to generate gene–cell count matrices. Filtered count matrices were used for downstream analysis with Seurat v5 in R (version 4.5.1(41)). Ambient RNA contamination was estimated and corrected using DecontX implemented in the celda package(42). Initial clustering information was provided to the algorithm. For cultured samples, default parameters were used. However, samples derived from freshly isolated retina contained substantial rod photoreceptor contamination, a frequent issue in retinal single-cell RNA sequencing(43). Therefore, the contamination prior parameter was increased (delta = 14) to improve contamination estimation. Sample-specific before- and after-decontamination bar plots were observed to monitor the effects of decontamination (shown in **Additional file 3**)(44). Doublets were estimated and removed via scDblFinder with default parameters, using preliminary cluster labels as input. Then, samples were loaded into Seurat (45) and extreme outliers, including low-quality cells and potential remaining multiplets, were filtered based on the number of detected genes and UMI counts. Because sequencing depth differed substantially between the data generated in the current study and data generated by Cowan and colleagues(26), we applied different filtering thresholds respectively. For in vivo samples, cells with fewer than 300 or more than 3,000 detected genes (nFeature_RNA), or fewer than 300 or more than 10,000 detected reads (nCount_RNA), were removed. For cultured samples, cells with fewer than 300 or more than 6,000 detected genes, or fewer than 300 or more than 25,000 detected reads, were excluded. In addition, cells with mitochondrial gene expression exceeding 10% (in vivo samples) or 15% (cultured samples, higher mitochondrial content expected due to conditions) were removed (46, 47).

Samples were then merged, and the RNA assay was split by sample ID, and SCTransform v2 normalization was applied (48) with mitochondrial percentage included as a covariate to be regressed out. Principal component analysis (PCA) was performed via *RunPCA*(), and the top 30 principal components were retained for downstream analyses. Harmony integration was applied(49) using the *IntegrateLayer()* function in Seurat v5 (method=”HarmonyIntegration”), which aligned shared cell types across datasets. Clustering was performed with *FindNeighbours*() and *FindClusters*() with resolution=0.6 to delineate the main retinal cell types. Clustered cells were projected into a low-dimensional space for visualization via RunUMAP(), and main cell types were identified based on the expression of canonical marker genes. Then, to address the partially remaining rod contamination(50, 51), cell type scores (CTS) were generated based on the expression of marker genes selected based on specificity and expression level (rod: *RHO, SAG, GNAT1, NRL, PDE6G, GNGT1*; Müller glia: *VIM, GLUL, RLBP1, SLC1A3*; cone: *ARR3, PDE6H, OPN1MW, CNGA3, GNAT2, GUCA1C*; bipolar cell: *PRKCA, GRM6, TRPM1, LRTM1, VSX1*; horizontal cell: *ONECUT1, ONECUT2, LHX1*; amacrine: *GAD1, GAD1, SLC6A9*; microglia: *AIF1, CX3CR1, P2RY12*). In addition to CTS, another score was calculated to roughly estimate the differentiation potential *of a cell (“DS”), based on the cumulative average* expression of *SOX2, ASCL1, NEUROG2,* and *NEUROD1*. Scores were generated and added to the object with *AddModuleScore*(). The threshold for CTS scores for separating cell types was empirically estimated for each cell type based on their expression distributions. CTS-based filtering first involved excluding cells with high CTS (CTS > 0.9) for more than one cell type and DS < 0.1 (for removing possible remaining doublets, multiplets, and highly contaminated cells). After this, cluster-specific iterative filtering was also applied, where subtle contamination was further removed, such as cells expressing rod photoreceptor genes in non-rod clusters (specific thresholds listed in **Table S5**). Because one of our main goals was identifying potentially differentiating cells, which might demonstrate a transient gene expression profile reflecting more than one cell type, only cells with a DS < 0.1 were filtered from the Müller glia cluster so as not to risk losing potentially transitioning cells.

After filtering, cells were re-normalized, re-integrated, and re-clustered; cluster annotation was finalized, and Müller clusters were subsetted for subsequent analysis, including re-normalization, re-integration, and re-clustering, which yielded 6 Müller cell subclusters. We listed the top significant markers of each Müller subcluster via *FindMarkers*() to gain insight about potential functional differences between subpopulations (DEGs listed in **Additional file 2A**). DEGs suggested that these subclusters reflected group- or treatment-related Müller glia states (such as quiescent, activated, metabolically active, stressed, or proliferative) rather than true Müller glia subtypes. To determine and visualize transcriptomic distance between Müller glia from different experimental groups, we calculated the Jensen-Shannon divergence (JSDiv) between pseudobulk expression profiles with the philentropy R package (version: 0.10.0). The JSDiv ranges from 0 to 1, where 0 indicates identical distributions, and thus can be used to quantify the similarity between expression profiles. We then computed the Jensen-Shannon distance (JSD), the square root of the JSDiv, which is a true metric suitable for further distance-based analyses. PERMANOVA test was applied on the JSDs to evaluate whether the average expression profiles (centroids) of the groups in multivariate space were significantly different. To ensure that the difference was not driven by differences in spread (i.e. how tightly samples clustered around their group average), we also tested beta dispersion. Visualizing after multidimensional scaling (MDS) confirmed that Müller glia from freshly isolated retinal tissue were distinct from both control and FGF-2-treated cultured conditions (JSD = 0.419, 0.412, respectively), whereas the two cultured conditions strongly overlapped (JSD = 0.115).

To identify DEGs between groups (freshly isolated, control culture, and FGF-2 treated culture), we applied the pseudobulk approach implemented in Seurat v5, aggregating counts across cells within each cluster to improve statistical power. The *FindMarkers*() function was utilized, specifying test.use = “DESeq2”. P-values were adjusted for multiple comparisons using the Benjamini-Hochberg method. We retained DEGs with adjusted p-values <0.05 (**Additional file 2B, C, D**). The percentage of cells expressing individual genes of interest (**Table 1** and **Table S2**) was calculated using the *Percent_Expressing*() function from the scCustomize package (v3.0.1) on log-normalized RNA expression values, with a default detection threshold of >0. To estimate the fraction of Müller glia exhibiting robust neurogenic transcriptional features, we quantified the percentage of cells across groups that expressed at least 3 of the following genes: *ASCL1, NEUROD1, OTX2, LHX4, LHX9, SOX2,* and also expressed *SOX9* and *VIM* at a more rigorous threshold of >0.5. For selected genes of interest (**Fig. 3I**), the proportion of cells expressing each gene within *SOX9*-positive Müller glia was quantified. Cells were classified as double-positive based on the same expression thresholds defined above (>0 for detection of the gene of interest; >0.5 for *SOX9* to ensure Müller glia identity), and percentages were computed per group for descriptive comparison.

### Statistical analysis of cell quantification

All statistical analyses of postmitotic cell counts obtained from microscopic images were performed using GraphPad Prism 9. Data normality was assessed with the Shapiro–Wilk test. Group comparisons were conducted using the Kruskal-Wallis test. Statistical significance was defined as *p* < 0.05 (**Additional file 1**).

## List of abbreviations

ACTA2: Actin alpha 2, smooth muscle
AIF1: Allograft inflammatory factor 1
ARR3: Arrestin 3
ASCL1: Achaete-scute homolog 1
BAM: Binary Alignment/Map
BCL: Base call file
BIO: 6-Bromoindirubin-3′-oxime
BrdU: 5-bromo-2′-deoxyuridine
BS: Blocking solution
BSA: Bovine serum albumin
CABP5: Calcium-binding protein 5
CALB1: Calbindin 1
CALB2: Calbindin 2/calretinin
CCN1: Cellular communication network factor 1
CD44: Cluster of differentiation 44
CNGA3: Cyclic nucleotide-gated channel alpha 3
COL1A2: Collagen type I alpha 2 chain
CRX: Cone-rod homeobox
CRYAA: Crystallin alpha A
CTS: Cell type score
CX3CR1: C-X3-C motif chemokine receptor 1
DAPI: 4′,6-diamidino-2-phenylindole
DEG: Differentially expressed gene
DIV: Days in vitro
DMEM/F-12: Dulbecco’s modified Eagle medium/nutrient mixture F-12
DNase: Deoxyribonuclease
DS: Differentiation score
DUSP6: Dual-specificity phosphatase 6
EdU: 5-ethynyl-2′-deoxyuridine
ELAVL3: ELAV-like RNA-binding protein 3
ELAVL4: ELAV-like RNA-binding protein 4
ETV1: ETS variant transcription factor 1
FGF-2: Fibroblast growth factor 2
GAD1: Glutamate decarboxylase 1
GAD2: Glutamate decarboxylase 2
GFAP: Glial fibrillary acidic protein
GJA1: Gap junction protein alpha 1
GLUL: Glutamate-ammonia ligase
GNAT1: G protein subunit alpha transducin 1
GNAT2: G protein subunit alpha transducin 2
GNGT1: G protein subunit gamma transducin 1
GRIK1: Glutamate ionotropic receptor kainate type subunit 1
GRM6: Glutamate metabotropic receptor 6
GSK-3: Glycogen synthase kinase 3
GUCA1C: Guanylate cyclase activator 1C
IBA1: Ionized calcium-binding adapter molecule 1
ID2: Inhibitor of DNA binding 2
JAK/STAT: Janus kinase/signal transducer and activator of transcription
JSD: Jensen–Shannon distance
JSDiv: Jensen–Shannon divergence
LGALS3: Galectin 3
LHX1: LIM homeobox 1
LHX4: LIM homeobox 4
LHX9: LIM homeobox 9
LRTM1: Leucine-rich repeats and transmembrane domains 1
MDS: Multidimensional scaling
MSG: Monosodium glutamate
mTOR: Mechanistic target of rapamycin
NEAT1: Nuclear paraspeckle assembly transcript 1
NeuN: Neuronal nuclei
NEUROD1: Neurogenic differentiation 1
NEUROG2: Neurogenin 2
NPVF: Neuropeptide VF precursor
NQO1: NAD(P)H quinone dehydrogenase 1
NR2E3: Nuclear receptor subfamily 2 group E member 3
NRL: Neural retina leucine zipper
OCT: Optimal cutting temperature compound
ONECUT1: One cut homeobox 1
ONECUT2: One cut homeobox 2
ONL: Outer nuclear layer
OPN1LW: Long-wave-sensitive opsin 1
OPN1MW: Medium-wave-sensitive opsin 1
OPN1SW: Short-wave-sensitive opsin 1
OTX2: Orthodenticle homeobox 2
P2RY12: Purinergic receptor
P2Y12 PAX6: Paired box 6
PB: Phosphate buffer
PBS: Phosphate-buffered saline
PBST: Phosphate-buffered saline with Tween-20
PCA: Principal component analysis
PCP2: Purkinje cell protein 2
PDE6G: Phosphodiesterase 6G
PDE6H: Phosphodiesterase 6H
PERMANOVA: Permutational multivariate analysis of variance
PFA: Paraformaldehyde
PKCα: Protein kinase C alpha
PROX1: Prospero homeobox 1
PTPRC: Protein tyrosine phosphatase receptor type C
RBPMS: RNA-binding protein with multiple splicing
RHO: Rhodopsin
RLBP1: Retinaldehyde-binding protein 1
RNA: Ribonucleic acid
RT: Room temperature
S100β: S100 calcium-binding protein beta
SAG: S-antigen/visual arrestin
SCGN: Secretagogin
scRNA-seq: Single-cell RNA sequencing
SEPTIN7: Septin 7
SLC1A3: Solute carrier family 1 member 3
SLC6A9: Solute carrier family 6 member 9
SOCS3: Suppressor of cytokine signaling 3
SOX2: SRY-box transcription factor 2
SOX9: SRY-box transcription factor 9
STAT3: Signal transducer and activator of transcription 3
TAGLN: Transgelin
TJP2: Tight junction protein 2
TRIS-HCl: Tris(hydroxymethyl)aminomethane hydrochloride
TRPM1: Transient receptor potential cation channel subfamily M member 1
TUBB3: Tubulin beta 3 class III
UMAP: Uniform manifold approximation and projection
UMI: Unique molecular identifier
VIM: Vimentin
VSX1: Visual system homeobox 1
VSX2: Visual system homeobox 2
Wnt: Wingless/integrated

## Declarations

### Ethics approval and consent to participate

All eyes were enucleated prior to cardiac arrest in accordance with the Declaration of Helsinki. All experimental protocols were approved by the National Ethics Committee (ETT TUKEB 34851-2/2018/EKU and ETT TUKEB IV/5645-1/2021/EKU).

### Consent for publication

Not applicable

### Availability of data and materials

The single-cell RNA-sequencing (scRNA-seq) data generated in this study have been deposited in the European Genome-phenome Archive (EGA) under data access number EGAD50000002223 and will be made available under controlled access upon acceptance of the manuscript. The code generated during this study has been deposited at https://github.com/SE-Retina/human-retina-muller-glia-proliferation-scRNAseq. Additional datasets supporting the conclusions of this article are included in the supplementary data files

## Funding

Forefront - Hungarian Research Excellence Programme KKP_21 138726 (DPM, LG, AS)

Hungarian Higher Education Institutional Excellence Programme, Neurology Thematic Programme of Semmelweis University TKP 2021 EGA384 25 (AS)

National Research, Development and Innovation Office grant 2025-3.1.1-ED-2025-00008 (AS)

Hungarian University Research Scholarship Programme of the Ministry for Culture and Innovation from the source of the National Research, Development and Innovation Fund 2025-2.1.1-EKÖP-2025-00014 (DPM)

## Authors’ contributions

DPM, AS, LG, and TT designed and conducted the experiments, analyzed the data, and interpreted the results. DPM, FK, and LG cultured human retinal explants. DPM and FK performed histological analysis on human retinal cultures. TT analyzed scRNA-Seq data. AC and ZZN coordinated human tissue donations. BG contributed to the flow cytometry experiment. SP and BG contributed to scRNA-Seq experiments. AS supervised the work. DPM, AS, and TT wrote the manuscript.

## Acknowledgements

The authors are grateful to Botond Roska and Cameron Cowan for their assistance with the scRNA-seq experiments.

