## Supplementary information for "Proliferative Capacity and Neural Lineage Commitment of Müller Glia in the Adult Human Retina"

**This file includes:**

Figures S1 to S10

Tables S1 to S5

**Other supporting materials for this manuscript include the following:**

### Supplementary figures and tables

#### Figure S1

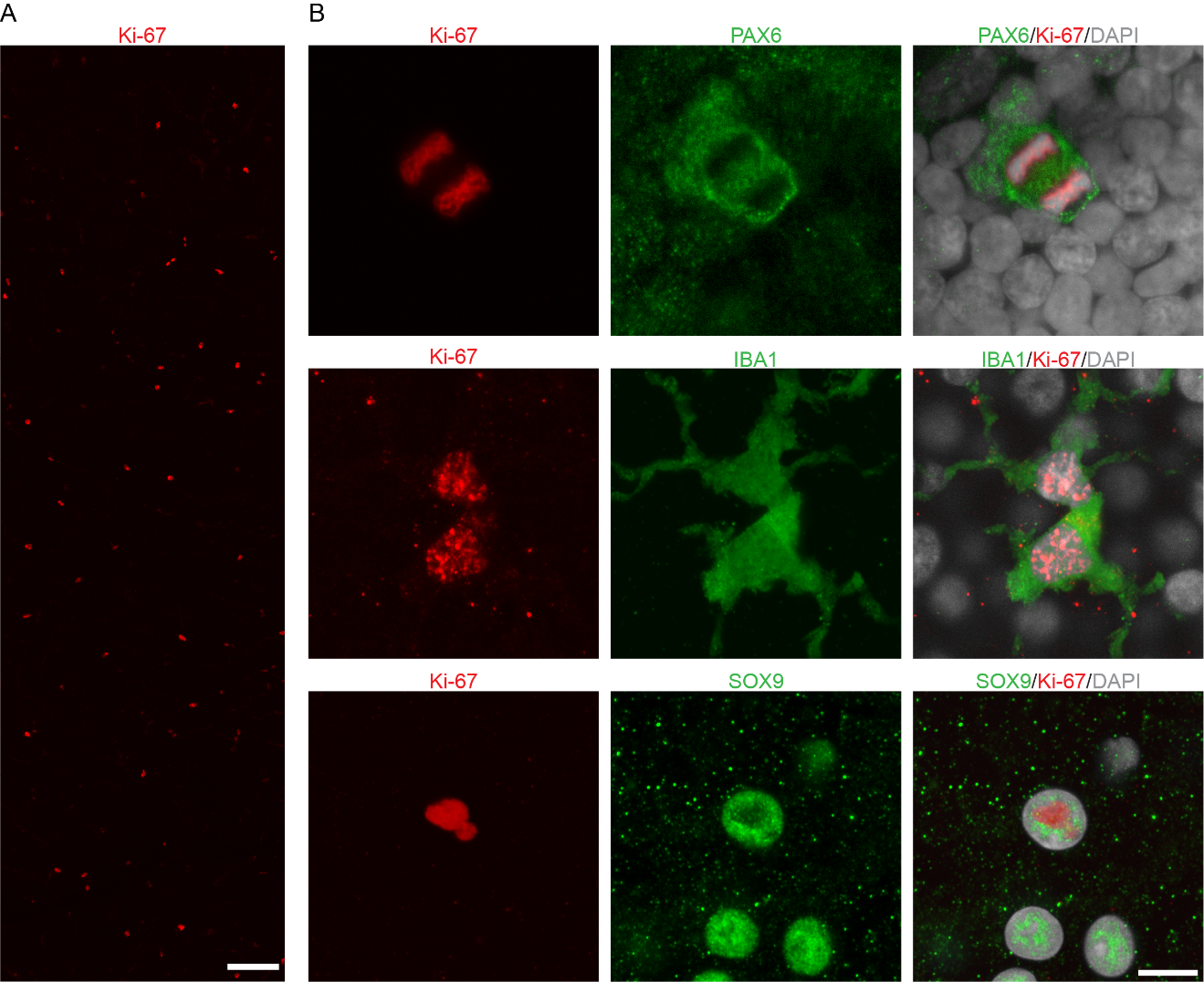

**Fig. S1** | **Proliferating cells in ex vivo human retina**

**(A)** Representative whole-mount image of ex vivo human retina fixed immediately after enucleation and immunolabeled for Ki-67 (red). A confocal image is shown as a z-projection reconstructed from optical sections spanning 27 μm. Scale bar: 100 µm.

**(B)** Representative whole-mount images of ex vivo human retina fixed immediately after enucleation, showing Ki-67–positive cells co-labeled with PAX6, IBA1, and SOX9. Confocal images are shown as z-projections reconstructed from optical sections spanning 1-3 μm. Scale bar: 10 µm.

#### Figure S2

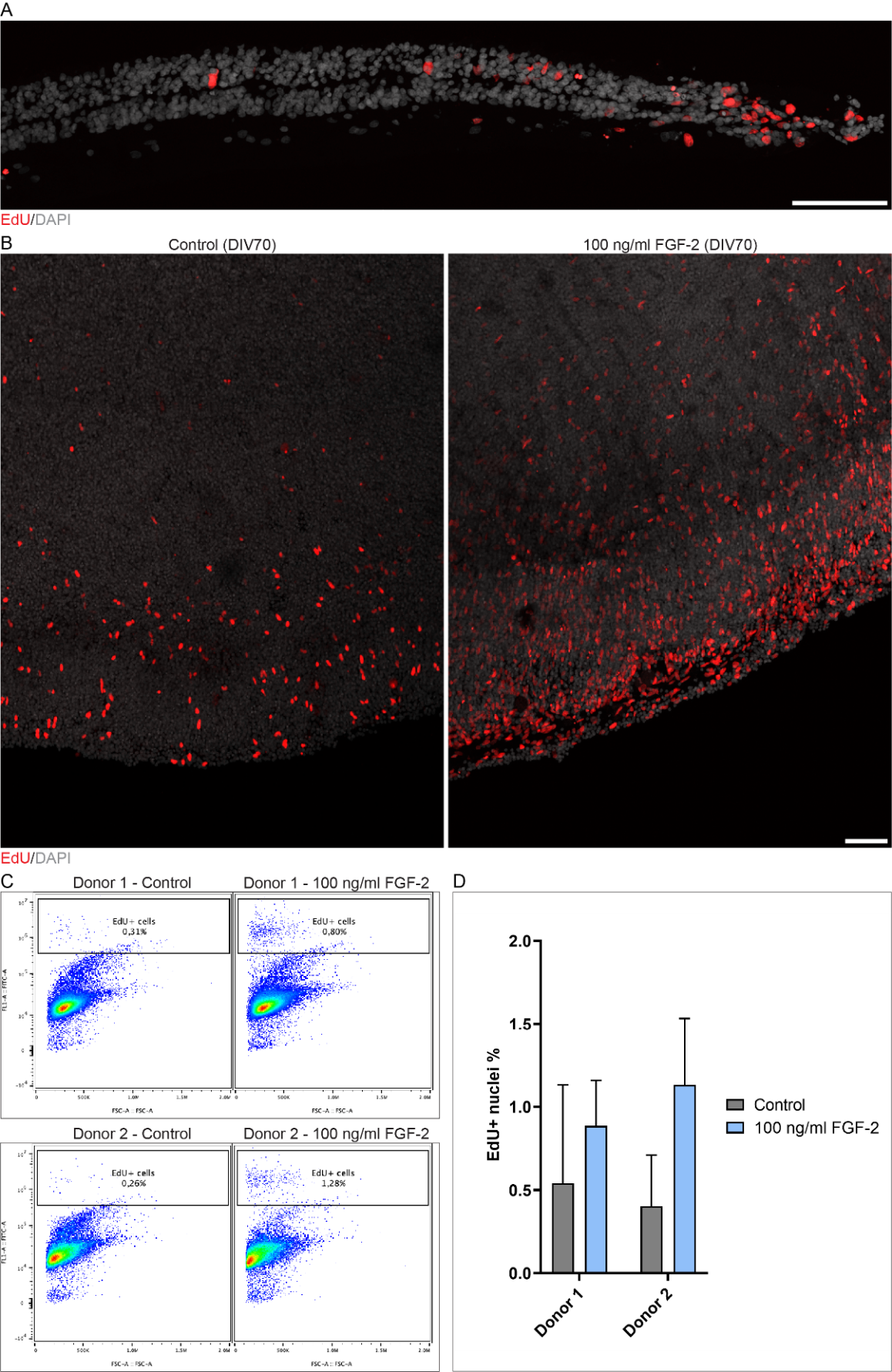

**Fig. S2 | Increased proliferation in cultured human retina**

**(A)** Representative image of the cultured human retina section edge with EdU (red) labeling demonstrating the effect of physical damage on proliferative activity (immunolabeled 10-week-old retinal section. Confocal image is shown as z-projection reconstructed from optical sections spanning 5 μm. Scale bar: 100 µm)

**(B)** Representative whole-mount image of EdU labeling in cultured human retina under control and high-dose FGF-2 conditions. EdU (red), DAPI (grey). Immunolabeled 10-week-old retinal whole-mount. Confocal images are shown as z-projections reconstructed from optical sections spanning 69-84 μm. Scale bar: 100 µm.

**(C)** Flow cytometry analysis of EdU incorporation in cultured human retina**.** After dissociation, the entire dissociated cell suspension was analyzed by flow cytometry. Hoechst was used to label nuclei, and FITC fluorescence was used to identify EdU-positive cells. Representative pseudocolor density plots of FSC-A versus FITC fluorescence are shown.

**(D)** Flow cytometric quantification of EdU+ cells in 10-week-old control cultures and cultures treated with 100 ng/ml FGF-2. Samples from two donors were analyzed (per donor: n = 4 control cultures and n = 4 FGF-2–treated cultures). For Donor 1, 61,738 Hoechst+ events were analyzed in control cultures and 73,564 in FGF-2–treated cultures; for Donor 2, 68,356 and 38,876 Hoechst+ events were analyzed, respectively. EdU+ fractions were compared between control and treated cultures using a two-tailed paired t-test. Data are shown as mean ± SD. Donor 1: 0.540 ± 0.593 (control) vs 0.885 ± 0.275 (FGF-2). Donor 2: 0.403 ± 0.307 (control) vs 1.133 ± 0.401 (FGF-2). Paired two-tailed t-test (matched by donor): t(1) = 2.795, P = 0.2188 (n = 2 donor pairs). Mean difference (FGF-2 − control) = 0.5376 ± 0.2721; 95% CI −1.907 to 2.982. For detailed statistics, see **Additional file 1.**

#### Figure S3

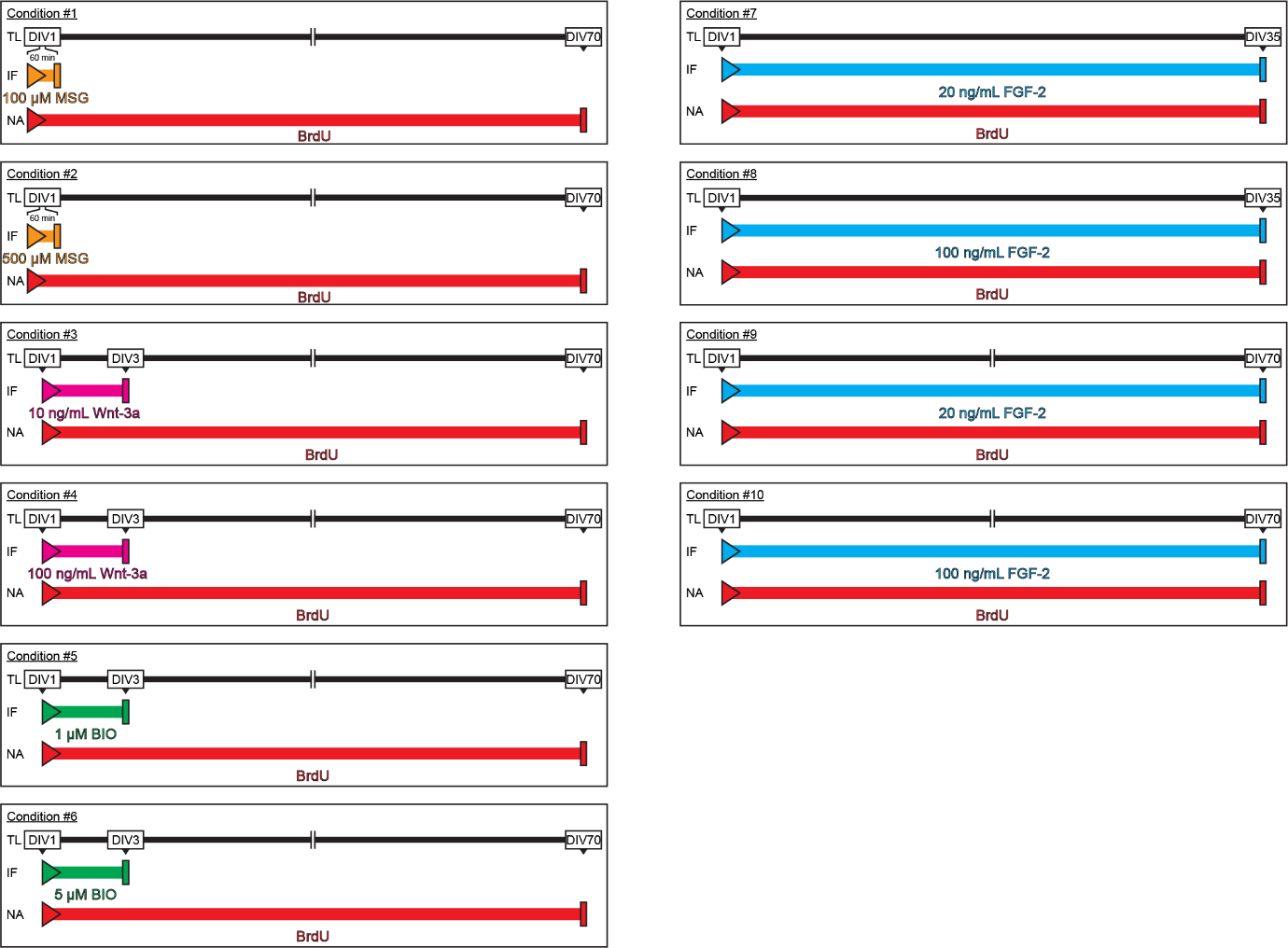

**Fig. S3 | Treatment conditions for quantification**

Schematics of the experimental conditions with MSG, Wnt-3a, BIO, and FGF-2 illustrating the treatment intervals and the culturing periods (DIV: days in vitro, TL: timeline, IF: induction factor, NA: nucleoside analog).

#### Figure S4

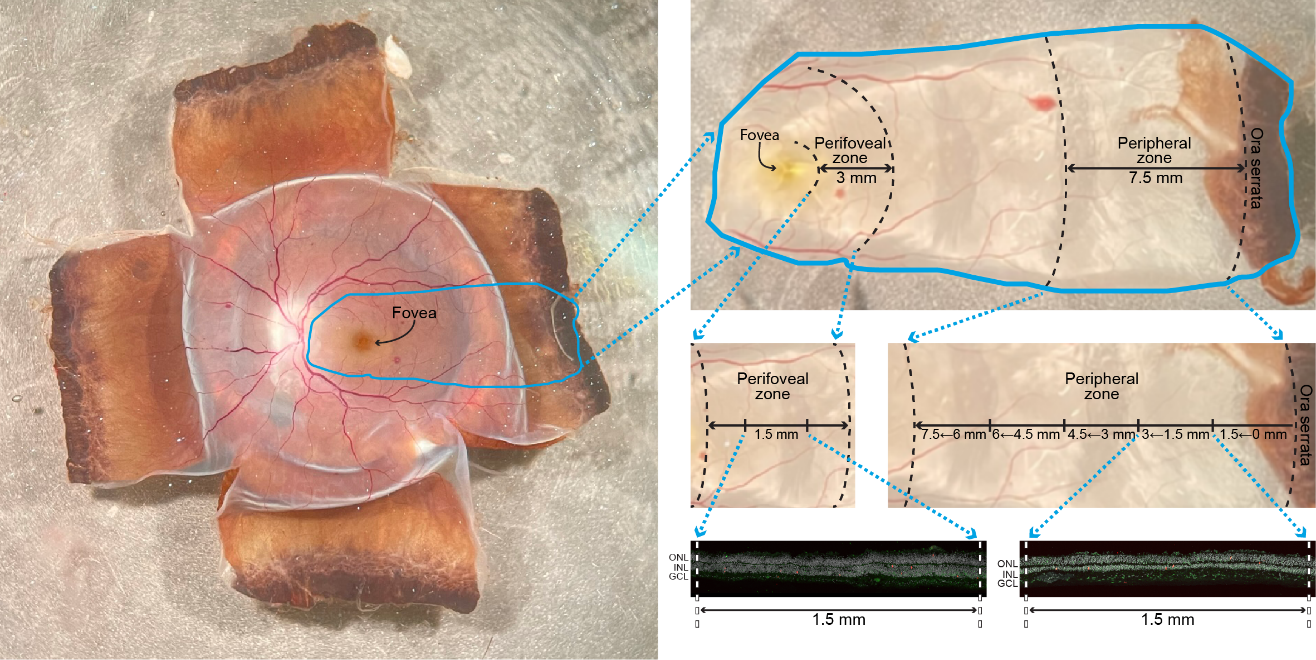

**Fig. S4 | Tissue sampling locations and sample sizes used for quantification**

Schematic of sampling locations in the whole retina used for perifoveal and peripheral quantifications, and the positions of the sections used for cell counting.

#### Figure S5

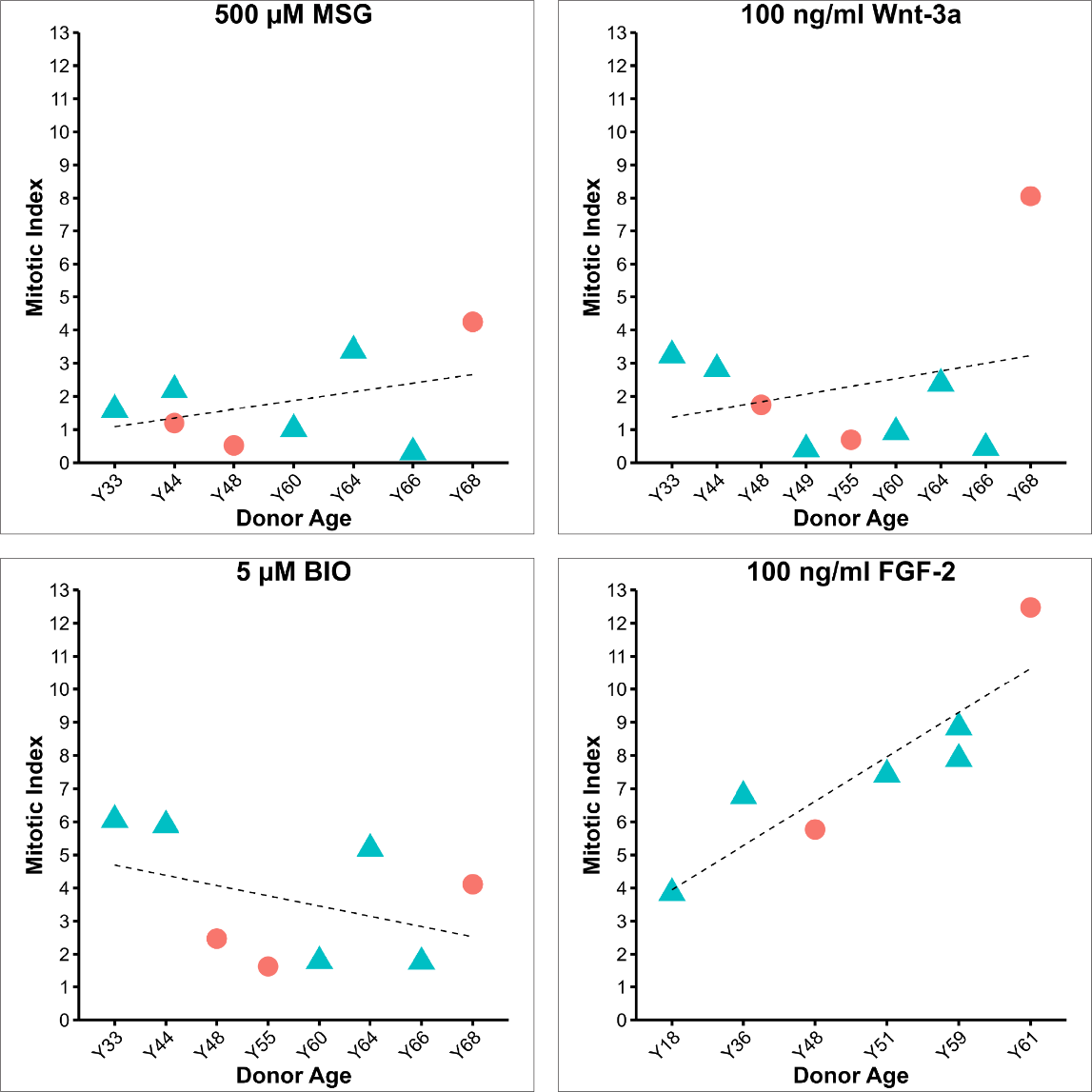

**Fig. S5 | Age and gender comparison of mitotic induction**

Mitotic responses to high-dose induction factors are shown by age group and gender. Mitotic index represents the normalized nucleoside analog–positive cell count under treatment relative to untreated controls. Each panel depicts responses to a distinct factor: 500 μM MSG (top left), 100 ng/ml Wnt-3a (top right), 5 μM BIO (bottom left), and 100 ng/ml FGF-2 (bottom right). Male (blue triangle) and female (red dot) donors are indicated. Data reflects high-dose treatments from the same peripheral dataset as **Fig. 1A**. Linear regression lines depict overall trends across donors. The numerical values for individual samples are listed in **Table S1**.

#### Figure S6

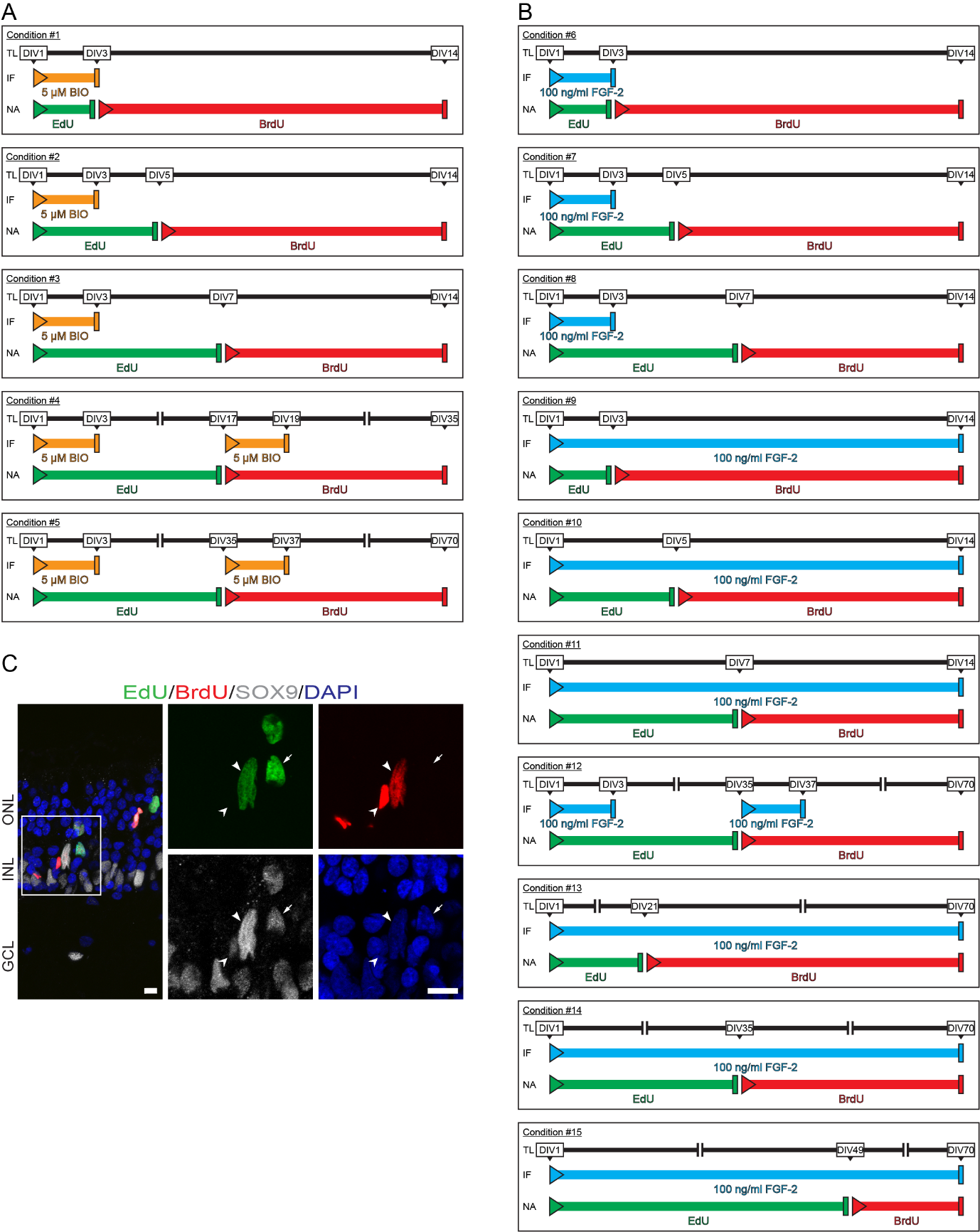

**Fig. S6 | Analysis of multiple divisions of Müller glia**

**(A)** Schematic of the experimental conditions using high-dose BIO, showing treatment intervals and the alternating time points for EdU and BrdU administration. (DIV: days in vitro; TL: timeline; IF: induction factor; NA: nucleoside analog)**.**

**(B)** Schematic of the experimental conditions using high-dose FGF-2, illustrating treatment intervals and the alternating time points for EdU and BrdU administration. (DIV: days in vitro; TL: timeline; IF: induction factor; NA: nucleoside analog).

**(C)** Representative image of cultured retina treated with high-dose FGF-2, showing EdU/BrdU/SOX9 triple-labeled cells (arrowhead), EdU/SOX9 double-labeled cells (arrow), and BrdU/SOX9 double-labeled cells (cut-out arrowhead). A confocal image is shown as a z-projection reconstructed from optical sections spanning 2.5 μm. Scale bars: 10 µm.

#### Figure S7

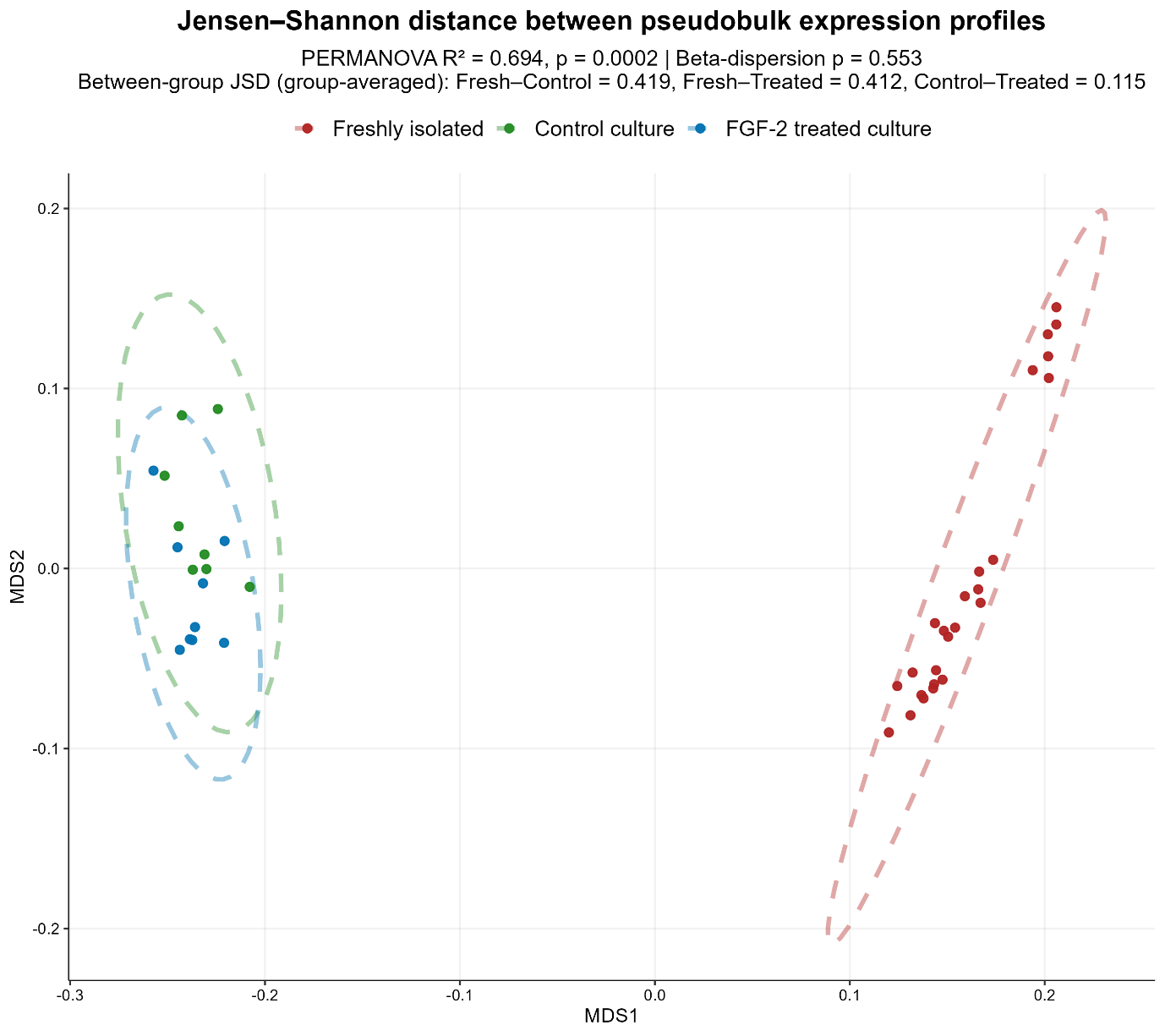

**Fig. S7 | Jensen-Shannon distances between expression profiles across** experimental **conditions**

Multidimensional scaling was performed on pairwise Jensen–Shannon distances computed between sample-level pseudobulk transcriptomic profiles (freshly isolated retina, control culture, and FGF-2-treated culture). Dashed ellipses denote 95% confidence regions around group centroids. Ordination of JSD distances showed clear separation between freshly isolated and cultured samples, while control and FGF-2-treated cultures exhibited substantial overlap, consistent with pairwise JSD distances. PERMANOVA on the JSD distance matrix also detected a significant overall effect of condition on centroid location in compositional transcriptomic space. Homogeneity of multivariate dispersion was confirmed, indicating that observed differences were not driven by unequal within-group variability.

#### Figure S8

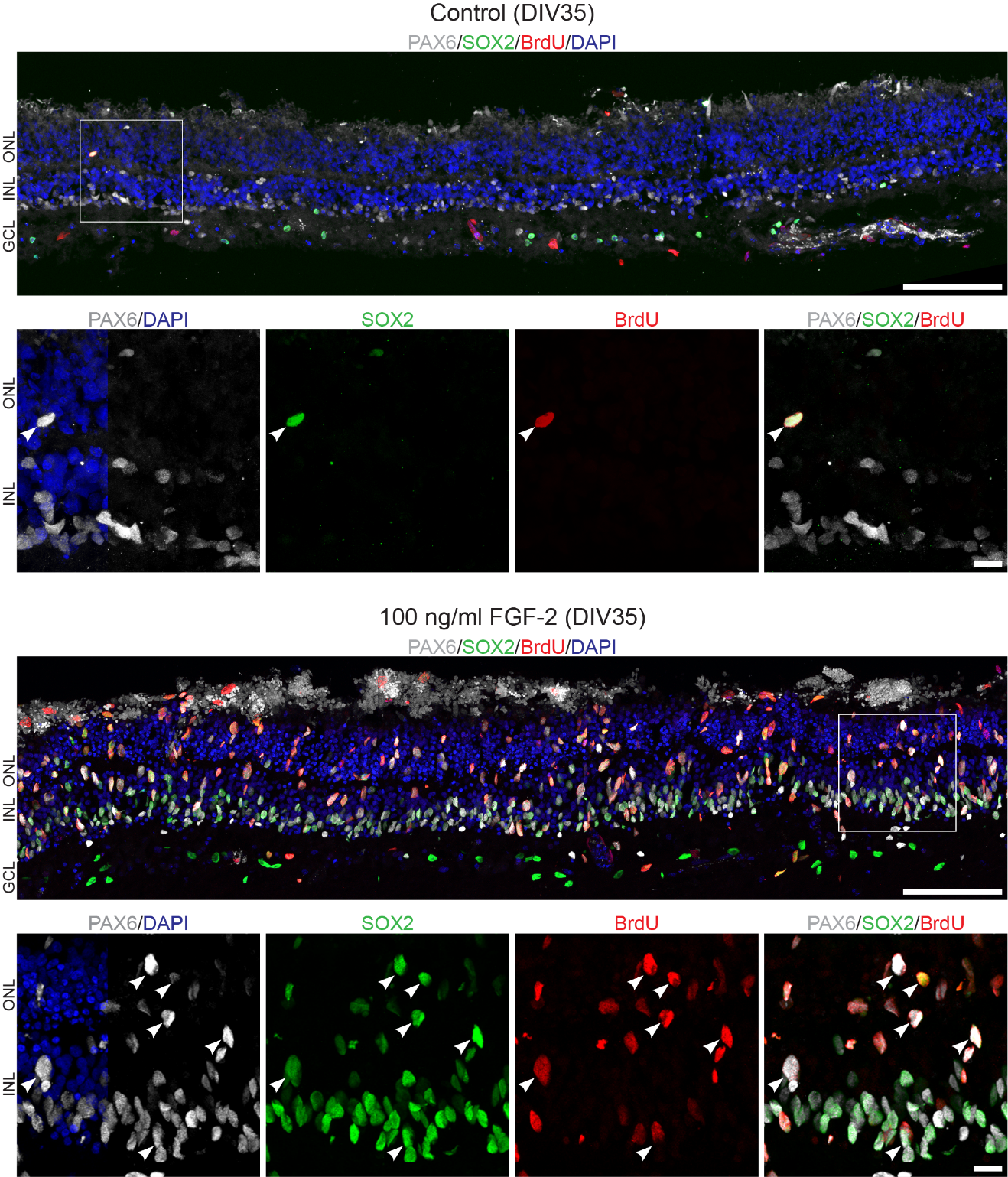

**Fig. S8 | Increased expression of progenitor markers in FGF-2-treated culture**

Representative images of cultured retina under control and high-dose FGF-2 conditions, showing increased SOX2 and PAX6 expression. Immunolabeled 5-week-old retinal sections, white rectangles indicate regions shown at higher magnification, arrowheads indicate examples of colocalizing cells, DIV: days in vitro. Confocal images are shown as z-projections reconstructed from optical sections spanning 14–15 µm, depending on the specimen. Overview image scale bar: 100 µm, higher magnification scale bar: 10 µm.

#### Figure S9

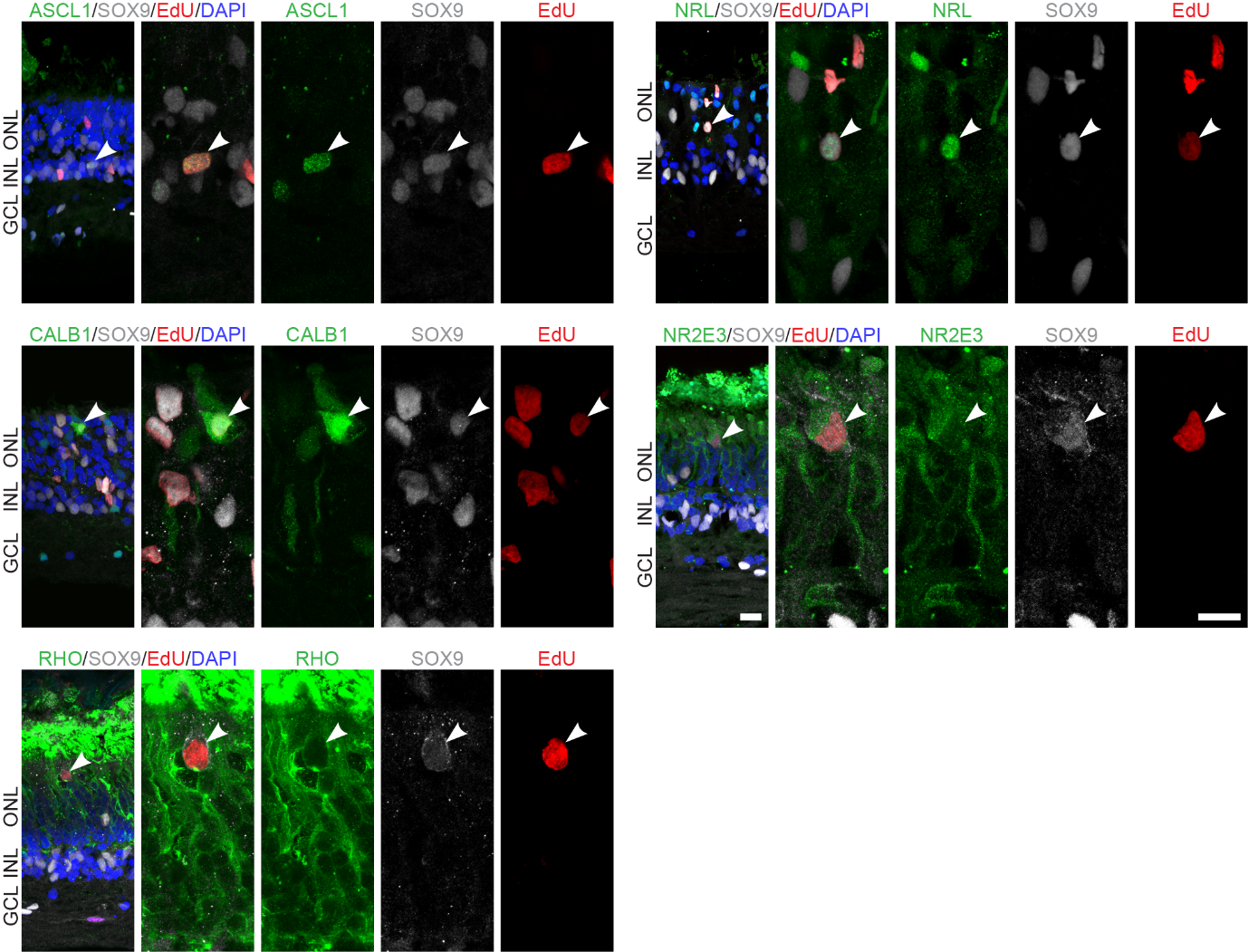

**Fig. S9 | Expression of neural markers in SOX9-positive postmitotic cells**

A more expanded and augmented representation of **Fig. 3I.** Postmitotic SOX9-positive cells co-express proneural- and neural markers in FGF-2-treated human retina**.** Representative confocal images showing triple-labeled immunofluorescence for the indicated markers (green: ASCL1, CALB1, NRL, NR2E3), SOX9 (gray), and EdU (red). Immunolabeled 10-week-old retinal sections, white arrowheads indicate the co-labeled cells. Confocal images are shown as z-projections reconstructed from optical sections spanning 1–3.5 µm, depending on the specimen. Scale bars: 10 µm.

#### Figure S10

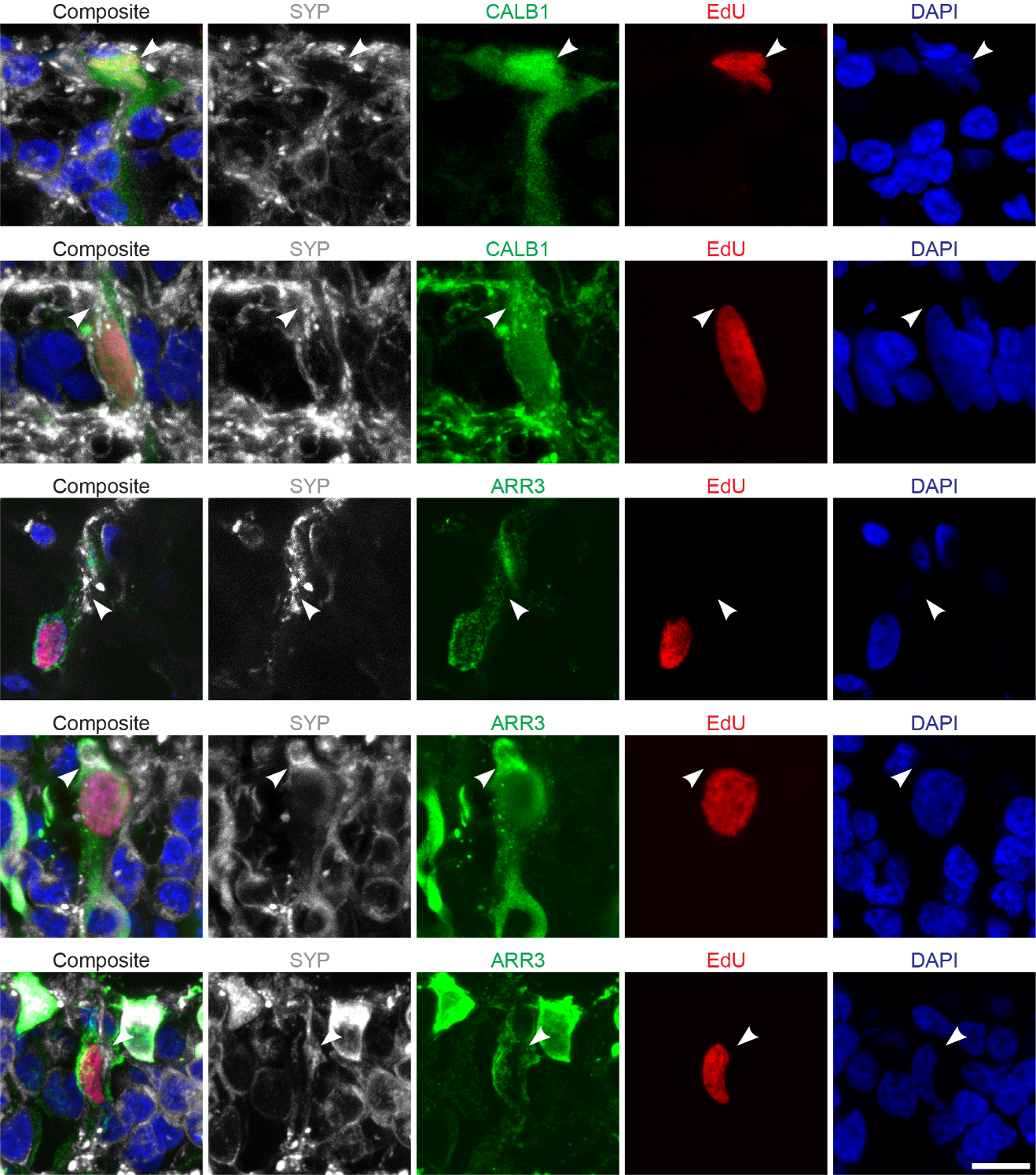

**Fig. S10** | **Synaptophysin expression in calbindin- and arrestin-3-positive postmitotic cells**

Representative confocal images showing Synaptophysin (SYN, gray) expression in cells labeled with the neural markers calbindin (CALB, green, top two rows) or arrestin-3 (ARR3; green, bottom three rows), together with EdU (red) and DAPI (blue) staining. Arrowheads indicate the cellular localization of Synaptophysin in EdU-positive postmitotic cells co-expressing respective neural markers. Confocal images are shown as z-projections reconstructed from optical sections spanning 1.5–3.5 µm, depending on the specimen. Scale bar: 10 µm.

#### Table S1

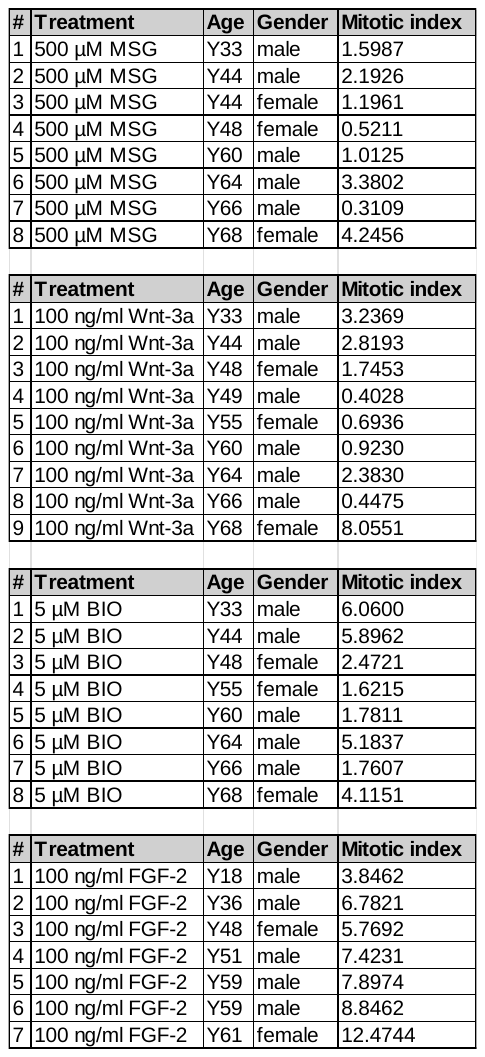

**Table S1 | Comparison of mitotic induction after high-dose treatments by age and gender (depicted and detailed in Fig. S5)**

#### Table S2

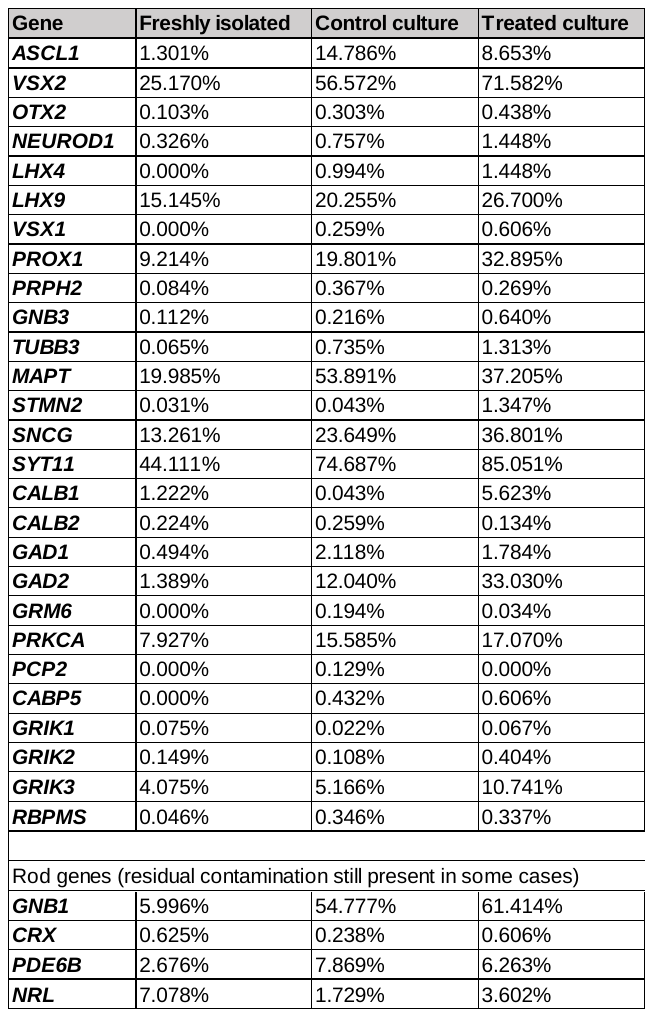

**Table S2 | Percentage of cells in the total Müller glia cluster expressing genes depicted on the proneural/neural dotplot (Fig. 3H)**

#### Table S3

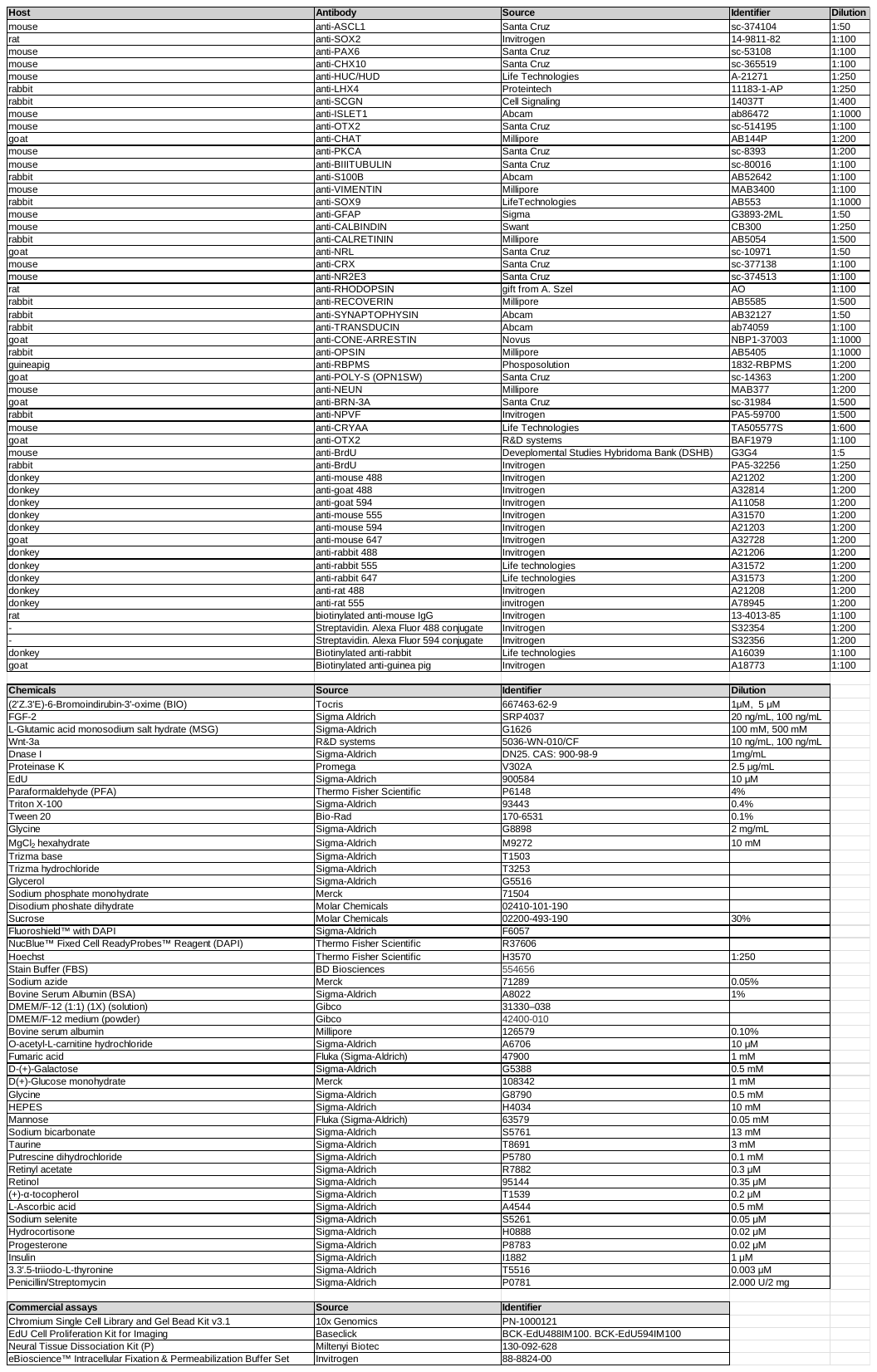

**Table S3 | List of antibodies, chemicals, and assays**

#### Table S4

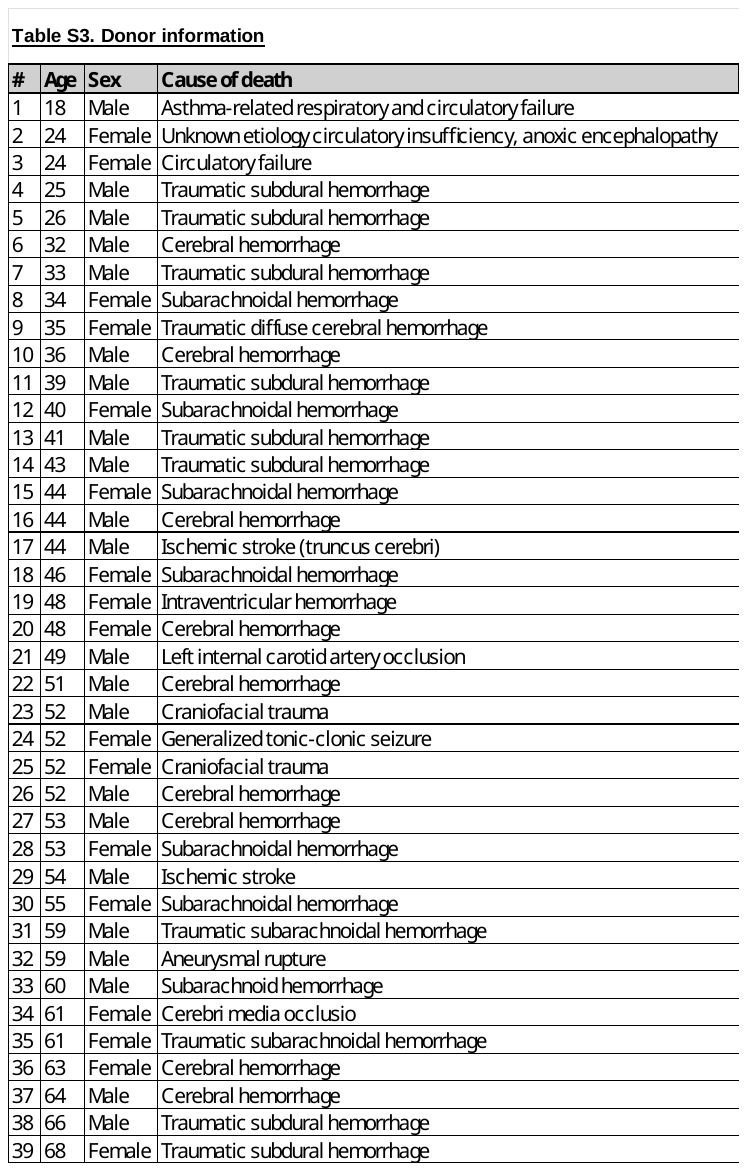

**Table S4 | Source of human tissue and donor information**

#### Table S5

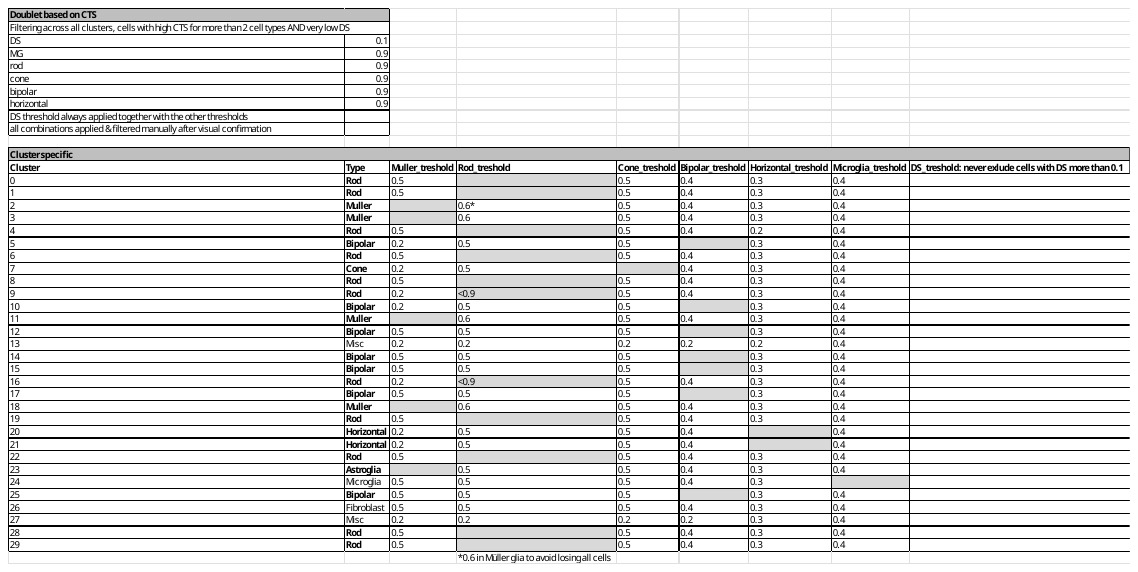

**Table S5 | Cell Type Score table**

#### Additional file 1 - Statistical data (not included in this version)

#### Additional file 2 – Marker genes and Differentially Expressed Genes (DEGs) (not included in this version)

#### Additional file 3 - DecontX filter comparisons (not included in this version)
